# Oncogenic RAS activity predicts response to chemotherapy and outcome in lung adenocarcinoma

**DOI:** 10.1101/2021.04.02.437896

**Authors:** Philip East, Gavin P. Kelly, Dhruva Biswas, Michaela Marani, David C. Hancock, Todd Creasy, Kris Sachsenmeier, Charles Swanton, on behalf of the TRACERx consortium, Sophie de Carné Trécesson, Julian Downward

## Abstract

Activating mutations in the driver oncogene *KRAS* occur in 32% of lung adenocarcinomas, leading to more aggressive disease and resistance to therapy in preclinical studies. However, the association between *KRAS* mutational status and patient outcome or response to treatment remains unclear, likely due to additional events modulating RAS pathways. To obtain a broader measure of RAS pathway activation beyond *KRAS* mutation only, we developed RAS84, a transcriptional signature optimised to capture RAS oncogenic activity in lung adenocarcinoma. Using RAS84 to classify lung cell lines, we show that RAS transcriptional activity outperforms *KRAS* mutation to predict resistance to chemotherapy drugs *in vitro*. We report that 84% of lung adenocarcinomas show clear transcriptional evidence of RAS oncogenic activation, falling into four groups characterised by coincident mutation of *STK11/LKB1*, *TP53* or *CDKN2A*. Given that 65% of these RAS pathway active tumours do not have *KRAS* mutations, we find that the classifications developed when considering only *KRAS* mutant tumours have significance in a much broader cohort of patients. Critically, patients in the highest RAS activity groups show adverse clinical outcome and reduced response to chemotherapy. The stratification of patients using gene expression patterns linked to oncogenic RAS signalling activity instead of genetic alterations in cancer genes could ultimately help clinical decision making.

## Introduction

The *RAS* oncogenes are mutated in close to 20% of all human cancers, acting as drivers of tumour formation and progression. Point mutations occur mainly in codons 12, 13 and 61 of the three isoforms *HRAS, KRAS* and *NRAS*, decreasing the GTPase activity of their encoded proteins and resulting in the accumulation of the GTP-bound, active conformation. *KRAS* is the most mutated *RAS* isoform, with particularly high prevalence in pancreatic ductal adenocarcinoma (88%), colorectal adenocarcinoma (50%) and lung adenocarcinoma (32%)^1^. The extensive literature describing the role of mutant *KRAS* in proliferation, survival, metabolism and motility supports its significant role in tumour aggressiveness, metastasis and resistance to chemotherapy^2–5^. However, there is a lack of consensus in published studies regarding the predictive value of *KRAS* mutations for patient outcome or response to treatment with chemotherapy^6–8^. *KRAS* mutants can also modulate the tumour microenvironment by regulating the expression of numerous cytokines^9^. Moreover, we have demonstrated that *KRAS* mutation promotes the expression of PD-L1, leading to immune evasion in models of human and mouse lung adenocarcinoma^10^. However, although it is clear that *KRAS* mutation does not preclude response to PD-1 immune checkpoint blockade^11, 12^, no consistent link between *KRAS* mutation and resistance to immunotherapy or PD-L1 expression has been shown in the clinic^13–16,11, 12^. Therefore, *KRAS* mutational status cannot be used as a predictive factor to select patients for specific therapy regimens^17^, with the exception of EGFR-targeted therapy, where *KRAS* mutations are negatively linked to response to EGFR inhibition in colorectal cancer^18^.

The stratification of patients uniquely on the mutational status of *KRAS* may have complicated the study of RAS mutants in large cohorts of patients. For instance, in The Cancer Genome Atlas (TCGA), 74% of the lung adenocarcinoma (LUAD) tumours are mutated in one or more genes from the broader RAS pathway, taken as running from receptor tyrosine kinases to ERK MAP kinases and phosphoinositide 3-kinases^19^. Here we propose a novel method of stratification based on RAS-regulated transcriptional activity which predicts outcome and response to treatment in lung adenocarcinoma and other solid cancers. We derived a gene expression signature, RAS84, and applied machine learning techniques to build a classifier to stratify patients according to the expression of RAS84 in their tumour. Using this method, we discovered that RAS transcriptional activity predicted clinical outcome in lung adenocarcinoma and several other solid cancers, where KRAS mutation alone did not. When applied to a cohort of chemotherapy-treated patients, our classifier predicted poor response in subjects with the highest RAS84 level of expression. We anticipate that the use of RAS84 to stratify patients will validate observations that were made in preclinical models of KRAS-mutant cancers, but not confirmed in clinical studies. Our method will offer the possibility to study the impact of oncogenic RAS activity in large cohorts of patients and may help to predict sensitivity to treatment associated with oncogenic RAS activity.

## Results

Other members of the RAS pathway signalling network in addition to KRAS can be altered and affect RAS pathway activity. To overcome this issue, we used RAS pathway transcriptional activity as a fingerprint of RAS oncogenic activity. To derive our meta-signature, we identified several RAS gene expression signatures previously established in different models from published data as well as our own hitherto unpublished data (Supplementary table 1). We selected studies performed in mouse models or human cell lines, where RAS activity was modulated using different methods, such as RNA interference or inhibition or over-expression of the mutant proteins, and where different isoforms and mutants were represented (KRAS mutants G12D, G12V, G12C, G12A, G13D, Q61H, HRAS mutant G12D). The datasets also represented several organs (lung, pancreas, colon, breast, kidney and prostate). To mitigate the possibility of confounding signal from tumour infiltrating immune cells, we removed all genes present in two immune signatures^20, 21^. We also assessed the overlap between the gene signatures. Although all were composed of RAS-target genes, we observed little commonality between the signatures (Supplementary Figure 1a).

### RAS84 construction using lung cell line expression data

Our initial goal was to measure RAS pathway activity in tumour cells. We therefore mapped the established signatures to lung cancer cell line data from the Broad Institute Cancer Cell Line Encyclopedia (CCLE)^22^ to determine which ones accurately measured oncogenic RAS activity in samples exempt of stroma and immune cells and where *KRAS* mutation is known to be a prevalent cancer driver. We first cleaned up the signatures by removing genes with low expression or variance across the cell lines (Supplementary Figure 1b). We removed cell lines with oncogenic RAS pathway mutations other than *KRAS* (*BRAF*, *EGFR*, *ERBB2*, *FGFR1*, *FGFR2*, *FGFR3*, *HRAS*, *JAK2*, *KIT*, *NRAS* & *RET*) from the analysis, since these mutations may drive RAS signalling and confound the analysis. For each signature, we clustered the filtered CCLE lung cancer cell line signature expression matrix into three groups. We named the clusters RAS-high and RAS-low according to the mean expression of the signature genes within the groups and categorised as “unclassified” the group of samples with intermediate mean expression (Supplementary Figure 1c). We used the distribution of *KRAS* mutations across the RAS-high and RAS-low clusters to assess the ability of the signature to capture RAS oncogenic activity (Figure 1a). We reasoned that the signatures measuring RAS oncogenic activity in the lung cell line dataset would show enrichment of *KRAS* mutations in the high group given its role in tumour development in the lung. To highlight the specificity of the signatures to measure RAS oncogenic activity, we also assessed a RAS addiction signature^23^ and several other oncogenic signalling pathway signatures^24, 25^. The RAS addiction signature was derived from KRAS mutant cell lines dependent on RAS signalling to maintain cell survival. This signature is therefore expected to capture RAS-dependency but not RAS oncogenic activity. We identified “RAS pathway”, “KRASG13D134” and “HRAS” as the best-performing signatures to enrich KRAS mutants in the RAS-high group (p-value < 1e-5) (Figure 1b). We refined these three signatures by selecting the genes driving the clustering of the RAS-high and RAS-low groups. We ran a differential gene analysis between these groups to identify signature genes upregulated in the RAS-high group of cells (FDR < 0.05) (Supplementary Figure 1d). From these genes, we constructed our meta RAS activity signature, RAS84, and tested it against the CCLE lung cancer cell line data (Figure 1c, Supplementary table 2). RAS84 successfully placed 36 out of 42 KRAS mutant lines into the RAS-high group, with six unclassified and none in RAS-low. When compared with other RAS and oncogenic signatures, RAS84 gave the most statistically robust separation of the KRAS mutant cell lines from the RAS-low group (Figure 1b and Supplementary Figure 1e). We ran a differential analysis to identify RAS-high dependent transcriptional changes when compared to RAS-low (2182 genes, fdr< 0.05, -1>LFC>1). We found ERK1 and ERK2 cascade and MAPK cascade GO terms (GO:0070371, p-value 4e-7; GO:0000165 p-value 9e-6) to be enriched in these genes (Supplementary table 3).

**Figure 1.**
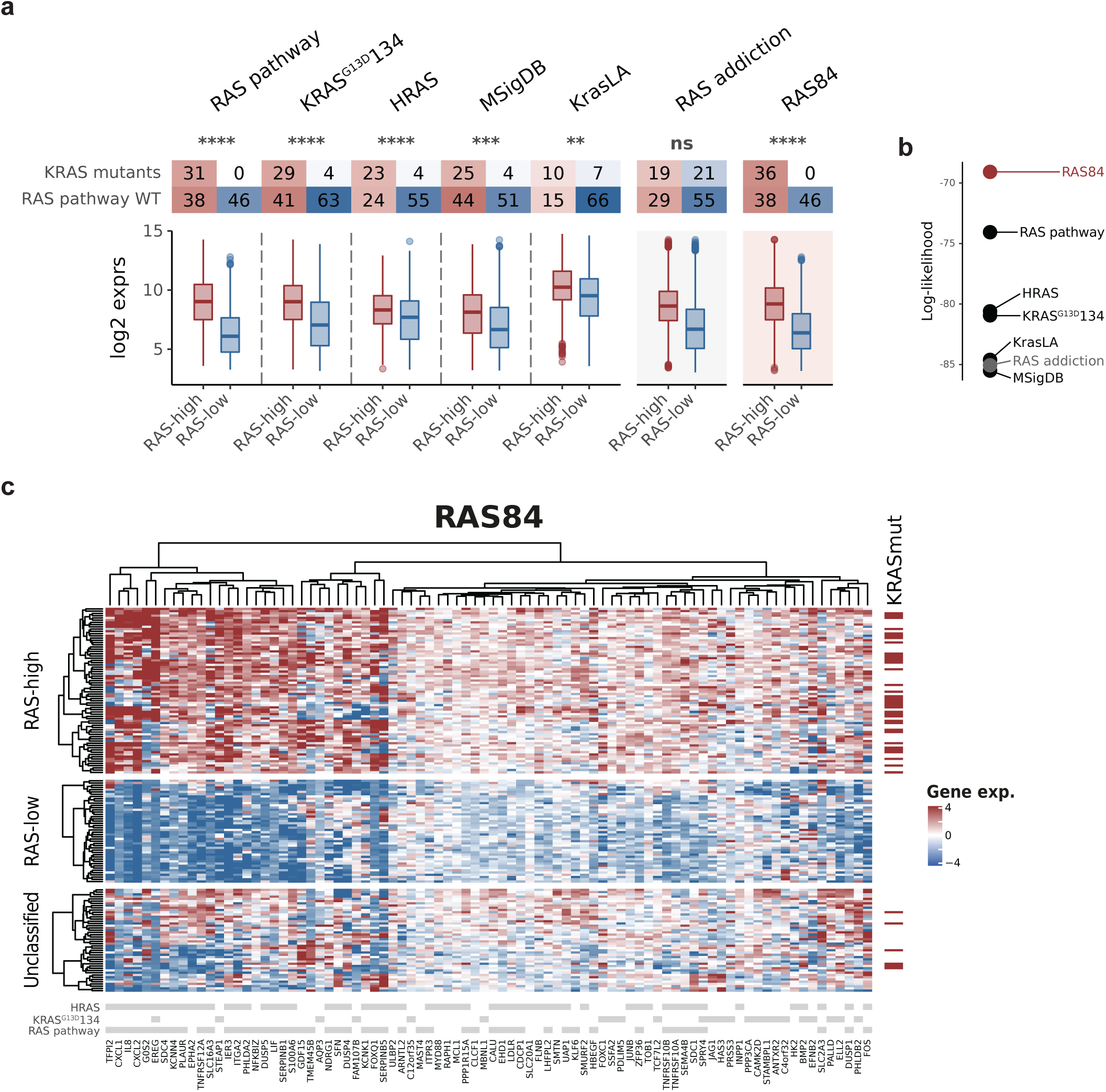
In vitro RAS signature derivation. **(a)** Contingency tables showing the number of RAS pathway wild type and KRAS mutant cell lines per RAS-high and RAS-low groups for each signa­ ture. RAS pathway wild type cell lines are those with no oncogenic mutation in any RAS pathway member. RAS-high cell counts are shown in red, RAS-low in blue. The boxplots show the RI distri­ butions for the RAS-high and low groups. RAS addiction is presented here as a control signature. (b) Log-likelihood values from a GLM fit (family= binomial) of KRAS mutation status across the three RAS activity groups for each of the signatures. This shows RAS84 to be the signature that per­ forming the best at segregating KRAS mutants across the RAS activity groups. (c) Heatmap show­ ing our RAS84 meta signature genes mapped to filter ed (see method) CCLE lung cell line data. Cell lines are shown as rows, genes as columns. Groupings of high, medium and low RAS activity are shown as separate clusters, KRAS mutational status is indicated in dark red on the right and parent signature gene membership is indicated in grey at the bottom of the map.

Using cell line gene expression and *KRAS* mutation data from lung cancer cell lines, we thus demonstrated the ability of RAS expression signatures to measure oncogenic RAS activity in a lung cancer context. We constructed a meta-signature from the best-performing signatures and demonstrated that it performed better than previous signatures at measuring RAS oncogenic activity by classifying KRAS mutant cell lines as RAS oncogenic signalling activated (RAS-high).

### RAS84 expression predicts drug sensitivity and resistance *in vitro*

To determine whether RAS84 expression was associated with anticancer drug response, we analysed drug sensitivity data obtained from the Genomics of Drug Sensitivity in Cancer project (GDSC) and The Cancer Therapeutics Response Portal (CTRP) in the context of RAS high and low CCLE cell lines. We identified drugs with differential drug responses across the two groups (GDSC fdr < 0.05, -1 > log2(delta IC50) > 1) (Figure 2a, Supplementary Figure 2a and Supplementary table 4-5). We tested for enriched drug target terms within the drugs showing differential response (hypergeometic fdr< 0.05_) (Figure 2b). We found RAG-high cell lines were sensitive to drugs targeting ERK MAPK and EGFR signalling and protein stability and degradation. We were encouraged to see sensitivity to ERK MAPK and EGFR signalling inhibition, confirming high dependence to RAS signalling in these cell lines. Conversely, we found RAG-high cell lines to be resistant to drugs targeting DNA replication, Mitosis and Chromatin histone acetylation. DNA replication and mitosis are common chemotherapy targets indicting that RAS84 activity is associated with chemotherapy resistance *in vitro*. We also tested for KRAS mutation (Figure 2c) and RAS pathway mutation dependent drug responses (Figure 2d). We found both mutant groups were sensitive to just three drugs targeting ERK MAPK signalling. We did not observe resistance to any drugs in these comparisons. This result shows RAS84 better captures RAS-driven drug response than KRAS mutation alone or wider RAS pathway mutants, highlighting the importance of our transcriptional approach.

**Figure 2.**
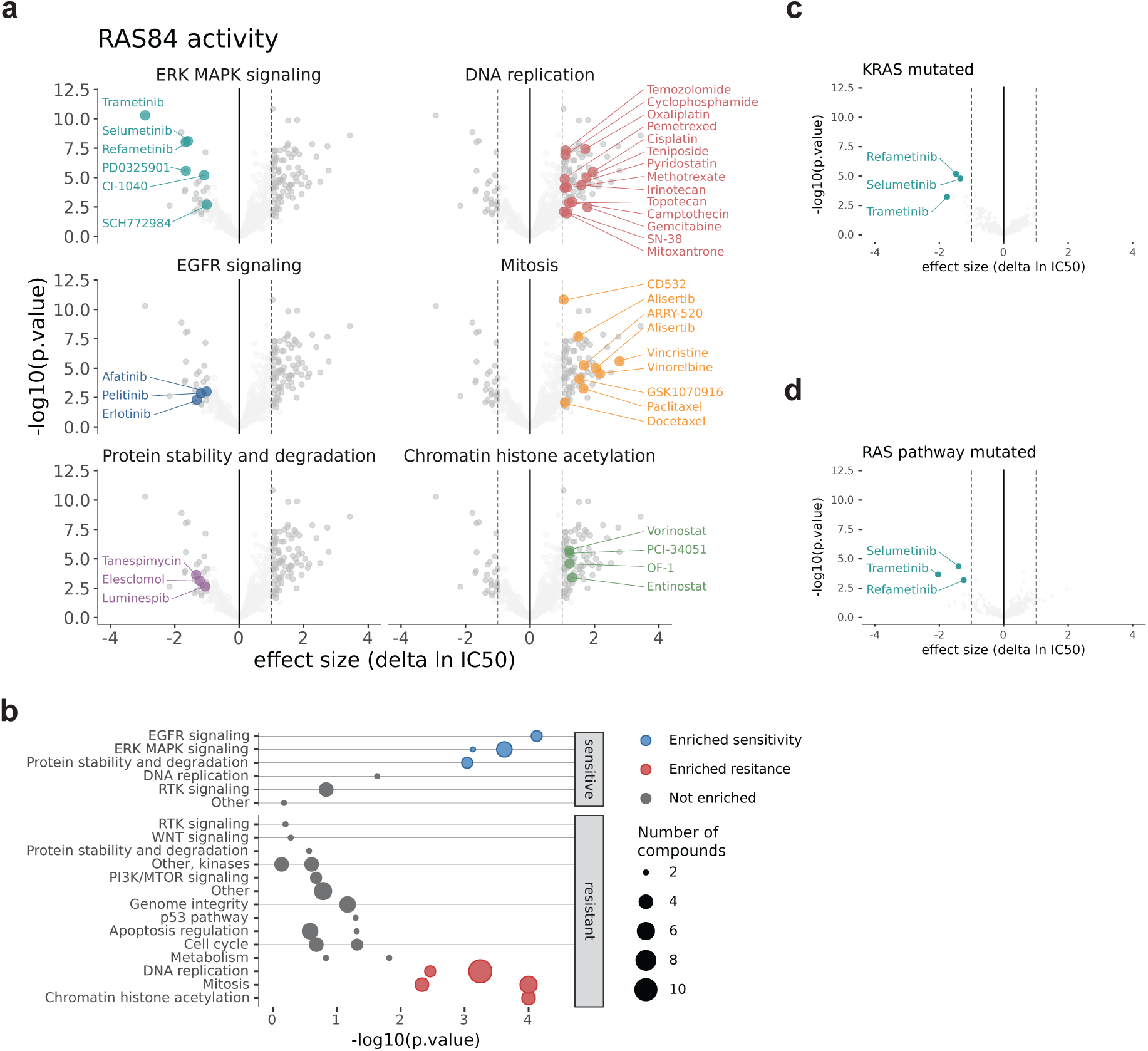
In vitro anti-cancer drug screen. **(a)** Volcano plots showing differences in IC50 values between RAS high and low CCLE cell lines. Drugs with enriched target annotations in the signifi­ cant sensitive and resistant groups are highlighted. Drugs with an absolute log2 fold change > 1 and fdr < 0.05 are shown in dark grey. Results from both GDSCl & 2 are shown. **(b)** Drug target annotation enrichment in sensitive and resistant drugs from GDSCl & 2 (fdr < 0.05) in the RAS high CCLE cell lines, determined by hypergeometic test. Target terms enriched in the sensitive drugs are shown in blue, the resistant in red. The number of drugs in each group is indicated by the size of the point. All test ed ta rgets are shown. **(c,d)** Volcano plots showing differences in IC50 values between KRAS mutant and wild type cell lines and RAS pathway mutat ed and wild type cell lines. Drugs with enriched target annotations in the significant sensitive and resistant groups are highlighted.

### RAS84 expression is associated with *KRAS* mutation in lung adenocarcinoma

To further validate RAS84 beyond cell lines, we applied it to clinical LUAD expression data from TCGA (512 samples)^26^. Given the increased heterogeneity in signature expression observed in patient tumour samples when compared to the cell line data, we explored the clustering of the patients beyond the three groups previously explained. We clustered the patients into five groups (Figure 3a) and found a low *KRAS* mutation count (6%) in the cluster with the lowest RAS84 expression (chi-square p-value 1.05e-08). The other clusters all had high levels of *KRAS* mutation, between 25 and 45% (Figure 3b). We regrouped the patients using the signatures described above and found RAS84 to better segregate *KRAS* mutations across the groups (Figure 3c). We labelled each RAS activity group (RAG) RAG-0, RAG-1, RAG-2, RAG-3 and RAG-4 ordered low to high by mean RAS84 expression. To ensure RAS84 expression was predominantly tumour-driven we looked at RAS84 expression in the stoma of five NSCLC samples^27^ (Supplementary Figure 3a) and found minimal expression. We assigned a RAS84-Index (RI) value to each patient, defined as the mean expression of the RAS84 genes. To further characterise these groups, we tested if any other reported genomic alteration^19^ exhibited a non-random distribution across the five clusters. We identified eight alterations enriched in one or more of the RAGs (Figure 3d) (chi-square, FDR < 0.05) (Supplementary Figure 3b) and used these cluster associated alterations to characterise each of the five RAGs (Figure 3a). *STK11*, *KEAP1*, *RB1*, *TP53*, *ATM* and *CDKN2A* (p-values: <2e-16, 6e-7, 3e-4, 5e-11, 5e-3 and 8e-4) are tumour suppressors whereas *EGFR* and *CTNNB1* (p-values: 2e-09 and 0.01) are proto-oncogenes. The underrepresentation of KRAS mutants characterised RAG-0, but interestingly this group contained a high number of tumour suppressor mutants (*KEAP1*, *RB1* and *TP53*) as well as *CTNNB1* mutants, which could explain how the tumours initiated. Due to the high level of p53 alterations (∼70%), we refer to these as P tumours. In addition to frequent *KRAS* mutations, many *EGFR* mutations also characterised RAG-2 and RAG-3, suggesting the genes driving these clusters reflected RAS pathway activation through upstream receptor tyrosine kinase (RTK) activation. *EGFR* and *KRAS* mutations are mutually exclusive^28, 29^. *TP53*, *STK11*/LKB1 and *CDKN2A* were identified as co-mutational partners of *KRAS* in NSCLC where the tumour suppressor gene mutations tend to be mutually exclusive^30^. Along with high *KRAS* mutation rates and RAS84 expression, RAG-1 was characterised by *STK11*/LKB1 mutations (KL tumours) and RAG-4 by *CDKN2A* mutations (KC tumours). *TP53* was frequently altered in several of the clusters, with RAG-3 having the highest rate of p53 alteration (hence KP tumours) after the RAS silent RAG-0. The RAG-2 cluster had modest levels of p53 (*TP53*) alteration (∼40%) and more beta catenin (*CTNNB1*) alterations than the other RAS active clusters, but still only about 5%. We refer to these as K tumours, as selective co-occurring mutations are not obvious. We validated the patterns of *KRAS*, *EGFR* and *TP53* mutations across our RAGs in an independent lung adenocarcinoma cohort of 87 patients (Seoul cohort, GSE40419) (chisq test p-values, *EGFR* 0.0002, *KRAS* 0.0014, *TP53* 0.1) (Figure 3e). Interestingly, we did not observe any significant association between specific KRAS amino acid mutational changes and RAG, suggesting that specific mutations do not drive RAS activity heterogeneity in distinct ways in the context of lung adenocarcinoma (Figure 3b).

**Figure 3.**
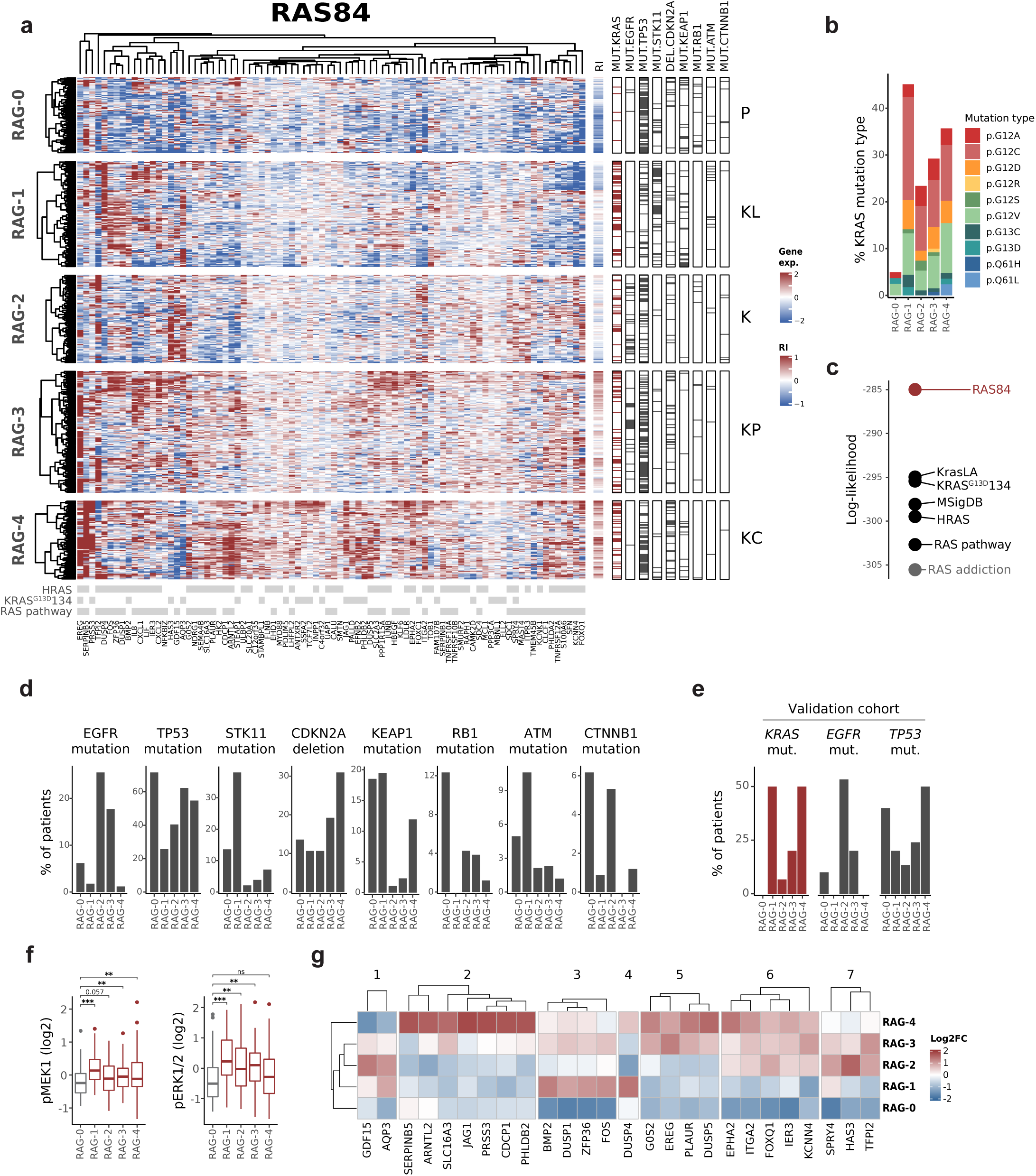
LUAD classification by RAS84. **(a)** Heatmap showing clustered RAS84 genes and TCGA LUAD cohort patients. Patients are shown as rows, genes as columns. Patients have been clustered into five RAS activity groups (RAGs) by hierarchical clustering using a ward.D2 agglomeration method. Aggregate RAS84-Index (RI) scores are shown to the right of the main heatmap. Genome variants with a significant non-random distribution across the RAGs are shown in the nine columns on the right (chi-square fdr < 0.05). These mutations are used to characterise the five clusters shown by the labels on the right (P, TP53; KL, KRAS/LKB1(STK11); K, KRAS; KP, KRAS/TP53; KC, KRAS/CDKN2A). KRAS mutants are shown in dark red. Parent signature membership is shown in grey at the bottom of the heatmap. **(b)** The percentage of KRAS mutations per RAG broken down by specific KRAS mutation type. **(c)** Log-likelihood values from a GLM fit (family=binomial) of KRAS mutation status across the five RAGs. **(d)** Bar plots showing the percentage of patients per RAG with EGFR, TP53, STK11 mutations, CDKN2A deletion, KEAP1, RB1, ATM and CTNNB1 mu-tations found to be significantly associated with any one RAG (fdr < 0.05). **(e)** EGFR, KRAS and TP53 mutation percentages found to be significantly associated with any one RAG from the Seoul cohort. **(f)** Boxplots showing The Cancer Protein Atlas (TCPA) RPPA MEK1 and ERK1/2 phosphoryl-ation level distributions across RAGs. Significance levels are shown compared to RAG-0 derived by linear model fit. **(g)** Heatmap showing variant mean RAS84 gene expression clusters across the five RAGs.

To determine if RAS84 expression reflected RAS-MAPK signalling activity we looked at ERK1/2 (T202, Y204) and MEK1 (S217, S221) phosphorylation levels within each RAG. We used The Cancer Proteome Atlas (TCPA) reverse-phase protein arrays (RPPA) data^31^ for 349 of the TCGA LUAD patients and found an increase in phosphorylation of one or both proteins in all RAGs when compared to RAG-0 (Figure 3f). At the expression level we found an enrichment of genes associated with the GO term ‘ERK1 and ERK2 cascade’ when comparing RAG-4 to RAG-0 (GO:0070371, p-value 0.0003) (Supplementary table 6). We also looked at the proliferation score distributions across the five groups and did not observe a correlation with RAS84 expression (Supplementary Figure 3c).

We were also interested in identifying which RAS84 genes were driving the RAGs. We focused on genes most variant across the clusters and, via correlation analysis (Supplementary Figure 3d), found seven gene clusters capable of discriminating the five RAGs (Figure 3g). RAG-4 was characterised by higher expression of genes in cluster 2 when compared to the other RAGs, RAG-3 by the expression of genes in cluster 5 but not 2, RAG-2 by the up-regulation of genes in clusters 6 but not 5 and to an extent the over-expression of cluster 1. RAG-1 patients could be identified by high expression of cluster 3 and low expression of cluster 6, RAG-0 by low expression of cluster 3. We found two clusters whose expression pattern across the five RAGs mirrored that seen in the enriched alterations.

Hence, we have demonstrated that RAS84 outperformed previous signatures in classifying KRAS mutant lung adenocarcinoma tumours as active for RAS-driven transcription. We identified five RAGs characterised by distinct associated mutational profiles and we showed RAS84 expression to be reflected at the protein level.

### RAS84 expression is clonal

The prognostic value of RAS84 makes it an attractive potential biomarker. A reliable biomarker should ideally, not be affected by the region of sampling and therefore not remain refractory to the intra-tumour heterogeneity observed in most cancers. Recent analyses of signatures derived for prognostication in lung cancer indicate that up to 70% of NSCLC tumours^32^ and 40% of LUAD tumours^33^ may be subject to sampling bias. To assess the intra-tumour heterogeneity of RAS activity in lung adenocarcinoma, we classified samples from the multi-region TRACERx cohort into our five RAS classification groups (102 samples from 41 patients)^34^. To classify the samples, we trained a support-vector machine (SVM) classifier using the TCGA LUAD classification results (see methods) and used it to assign RAG labels to the TRACERx samples (Figure 4a). This classifier will allow the stratification of new patient samples outside of cohort datasets enabling the clinical application of RAS84.

**Figure 4.**
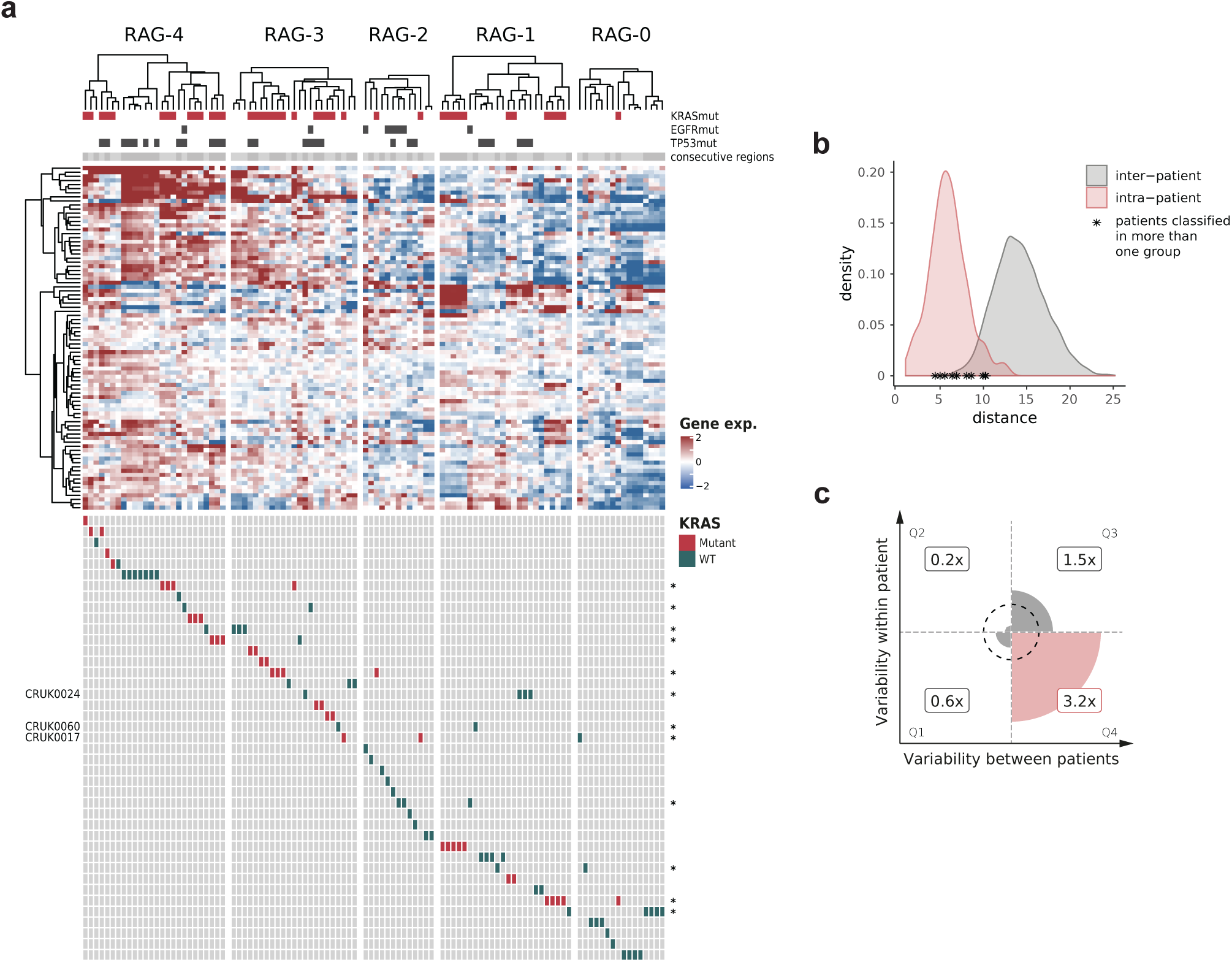
Intra-tumour RAS84 heterogeneity. **(a)** Heatmap showing RAS84 gene expression across the TRACERx multi-region cohort. The samples have been classified into the five RAGs using an SVM classifier. Contiguous samples from the same patient are indicated by the grey shades just at the top of the heatmap. Gene mutation status is indicated at the top, KRAS in red, EGFR and TP53 **in** grey. Per-patient regions are indicated by the rows at the bottom of the heatmap. KRAS mutants are shown in red, wild type in green. Patients with regions spanning multiple RAGs are indicated with an asterisk. Identifiers are given on the left in the three cases where regions did not fall into adjacent RAGs. (b) Intra- (orange) and inter-tumour (grey) sample Euclidian distant dis­ tributions. The maximum intra-patient distance for patients with samples spanning different RAGs a re indicated with an asterisk. (c) Plot showing the enrichment of RAS84 genes in genes previously classifie d as low for intra-tum our expression variance and high for inter-tumour expression variance in the TRACERx lung adenocarcinoma cohort. This group is represented by quadrant 4 on the plot showing a 3.2x fold enrichment of RAS84 genes relative to all expressed genes (Fisher’s exact p-valu e 9 .06e-6).

Of the 41 patients, 28 (68%) had multi-region RNA-seq gene expression data available. Of these 28, 16 (57%) patients had all regions falling within the same RAG, 13 of which clustered contiguously (Figure 4a). Twelve (43%) patients had regions that spanned RAGs suggesting a degree of RAS activity heterogeneity in some patients. All but three of these patients (CRUK0017, CRUK0024 and CRUK0060) span neighbouring RAGs indicating a degree of relatedness in RAS activity across the tumour. CRUK0017 spanned three RAGs R1:RAG-0; R2:RAG-3; R4:RAG-2. This patient is reported as sub-clonal for *KRAS* mutation by TRACERx (PhyloCCF R1:0; R2:0.84; R4:0.65). Interestingly, the *KRAS* non-mutated region (R1) fell into RAG-0. We observed that the intra-tumour distances for RAS84, within these group-spanning tumours, were still small when compared to the inter-tumour distance distribution (Figure 4b).

We assessed the intra- and inter-tumour expression variance of RAS84 genes by comparing them to gene-sets previously annotated for expression heterogeneity in the TRACERx lung adenocarcinoma cohort. Biswas and colleagues classified all expressed genes into four groups depending on their intra- and inter-tumour expression variance^33^. We found a significant 3.2 fold enrichment of RAS84 genes in the low intra-tumour, high inter-tumour expression variance group (Fisher’s exact p-value 9.06e-6) (Figure 4c, Supplementary table 7). This shows RAS84 genes tended to be enriched for genes robust to sampling bias.

Altogether, these data demonstrate that RAS activity is predominantly clonally expressed in lung adenocarcinoma, likely reflecting the oncogenic driving capability of the RAS pathway.

### RAS84 predicts survival and response to chemotherapy in lung adenocarcinoma patients

RAS oncogenic activity promotes tumour progression and metastasis but the mutational status of *KRAS* is not reliably associated with outcome^6, 7, 35^ (Figure 5a). To determine whether RAS84 had prognostic value in lung adenocarcinoma, we ran a univariate Cox proportional-hazards analysis comparing overall survival across the TCGA LUAD RAGs (n=493, 265 stage I, 117 stage II, 79 stage III and 25 stage IV). We found RAG-4 to be significantly associated with negative outcome when compared to RAG-0 (Figure 5b-c, Supplementary Figure 4a-f). We also fitted a univariate Cox proportional-hazards regression model to the RI values. We found a significant positive association with outcome, showing increased RAS84 expression was a predictor of poor overall survival (coxph HR 2, p-value 0.00042). To visualise the ability of RI to predict outcome we used the model to predict survival time given a two-fold increase or decrease in RI values (Figure 5d). Since we observed a slight over-representation of stage III tumours and an under-representation of stage I tumours in RAG-4 (Supplementary Figure 4g-h) we confirmed these findings in a multivariate Cox proportional-hazards analysis in an independent lung adenocarcinoma cohort of 103 patients (60 stage I, 19 stage II and 24 stage III) (Uppsala cohort, GSE81089). We first clustered the patients into five groups as previously described and ran the multivariate analysis across the RAGs and RI values including TNM stage, World Health Organization (WHO) performance status, smoking history, sex and age in the model. We found RAG to be a significant predictor of outcome (ANOVA LRT p-value 0.032) and specifically RAG-4, RAG-3 and RI to be significantly associated with poor outcome (Figure 5e-h, Supplementary figure 4i-j). We repeated these multivariate analyses with patient clusters derived using the signatures previously described (Figure 1). We found RAS84 to be the only signature significantly predicting outcome (Figure 5i, Supplementary table 8). This shows RAS84 had prognostic qualities in early stage lung adenocarcinoma beyond known clinical predictors and suggests RAS activity promotes tumour progression in human lung adenocarcinoma.

**Figure 5.**
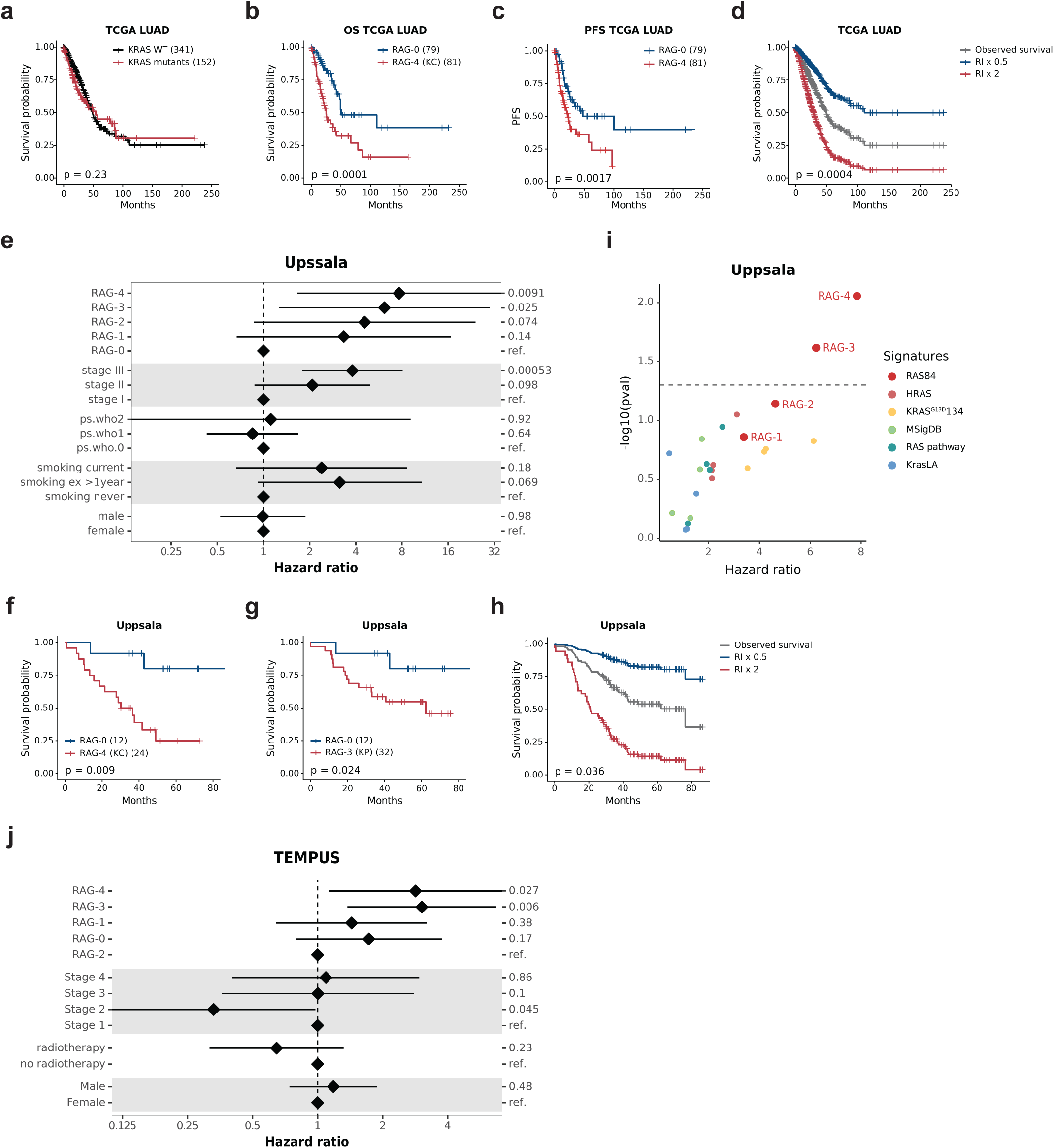
RAS84 predicts survival in lung adenocarcinoma. **(a)** Kaplan-Meier plot showing over-all survival data from the TCGA LUAD cohort for patients stratified by KRAS mutation (coxph p-val-ue 0.23), the number of patients per group in indicated in brackets. **(b)** Kaplan-Meier plot showing overall survival data from the TCGA LUAD cohort for patients from RAG-4 and RAG-0 (coxph p-value 0.0001), the number of patients per group in indicated in brackets. **(c)** Kaplan-Meier plot showing progression-free survival data from the TCGA LUAD cohort for patients from RAG-4 and RAG-0 (coxph p-value 0.0017), the number of patients per group in indicated in brackets. d, Kaplan-Meier plot showing a Cox proportional-hazards regression fit of TCGA LUAD survival data to RI values. The grey survival curve shows the observed data. The blue survival curve is the predicted survival if the RI values decreased 2-fold. The red curve are the predicted survival values if the RI were to increase 2-fold (coxph p-value 0.0004). e, Forest plot showing results from a multivariate Cox proportional-hazards analysis of the Uppsala lung adenocarcinoma cohort (n= 103 patients). RAG along with TNM stage, World Health Organization (WHO) performance status, smoking history, gender and age were tested. RAG-3 and 4 were significant after multivariate correction (coxph p-value RAG-4 0.0088, RAG-3 0.024), hazard ratios and 5 and 95% confidence intervals are shown on a natural log scale. f, Kaplan-Meier plot showing overall survival data from the Uppsala cohort for patients from RAG-4 and RAG-0 (multivariate coxph p-value 0.0088), the number of patients per group in indicated in brackets. g, Kaplan-Meier plot showing overall survival data from the Uppsala cohort for patients from RAG-3 and RAG-0 (multivariate coxph p-value 0.024), the number of patients per group in indicated in brackets. h, Kaplan-Meier plot showing a Cox proportional-hazards regression fit of Uppsala survival data to RI values. The grey survival curve shows the observed data. The blue survival curve is the predicted survival if the RI values decreased 2-fold. The red curve are the predicted survival values if the RI were to increase 2-fold (coxph p-value 0.036). i, Multivariate p-values and hazard ratios plotted for RAGs derived from RAS84 and the other RAS signatures. The p-values are plotted on a -log10 scale (coxph p-value RAS84 RAG-4 0.0088, RAS84 RAG-3 0.024). j, Forest plot showing results from a multivariate Cox proportional-hazards analysis of PFS after chemotherapy in the TEMPUS lung adenocarcinoma cohort (n= 100 patients). RAG high and RAG_0 are compared to RAG medium. Tumour stage, whether the patient received radiotherapy or not and sex were also tested (coxph p-value RAG-high p-value 0.0018). Hazard ratios and 5 and 95% confidence intervals are shown on a natural log scale.

Given that we observed a chemotherapy drug resistance phenotype *in vitro,* we ran a PFS multivariate Cox proportional-hazards analysis using the TEMPUS cohort of adenocarcinoma patients (n=94, 5 Stage I, 17 Stage II, 31 Stage III and 41 Stage IV). We selected patients who had received first line chemotherapy treatment and constructed PFS intervals using patient records (see methods). We classified the patient tumours using associated RNA-Seq data and our SVM classifier. We modelled PFS with RAG labels along with stage, the administration of radiotherapy, age and sex covariates. We found RAG to be a significant predictor of PFS after chemotherapy (ANOVA LRT p-value 0.043). Specifically, patients in RAG-3 and RAG-4 had a poor response when compared with RAG-2 (p-value 0.006, 0.027; HR 3.04, 2.84) (Figure 5j). We also ran the same multivariate analysis testing KRAS mutation as a predictor of PFS. As previously shown^7, 8^, KRAS mutation did not predict response to chemotherapy (Supplementary figure 4k). This result shows the potential of RAS transcriptional activity to predict response to chemotherapy where KRAS mutation status alone does not.

We thus demonstrate the prognostic value of RAG classification and RI quantification in 500+ lung adenocarcinoma patients from two independent cohorts, benchmarked against the failure of KRAS mutational status or previous RAS signatures to predict patient outcomes. We also show RAG classification as a predictor of response to chemotherapy, thus demonstrating that RAS84 adds value to current clinical risk factors and response biomarkers.

### RAS84 predicts RAS-MAPK activity across cancer types

The degree to which RAS activity is important in tumourigenesis and cancer progression varies across different tissues^19^, with some cancers known to be driven largely by RAS mutations (e.g. pancreatic, colorectal and lung cancers^36, 37^) and others not (e.g. uveal melanoma^38^, glioblastoma^39^, kidney cancer^40^). To determine how RAS84 varied across cancer types, we quantified it against all 32 TCGA solid cancers in a pan-cancer analysis. To compare samples across cancers, we calculated an RI value for each sample (Figure 6a). We identified two distinct cancer populations from the distribution of mean RI values per-cancer (Figure 6b, Supplementary Figure 5a). We found four of the top five RAS mutated cancers known to be RAS-driven (RAS mutation frequency: pancreatic adenocarcinoma (PAAD) 71%, colon adenocarcinoma (COAD) 50%, rectum adenocarcinoma (READ) 49% and lung adenocarcinoma (LUAD) 31%) in the highly RAS active group (Supplementary figure 5b). We also found *KRAS* mutation to be over-represented within this group (hypergeometric p-value < 2e-16). The other cancers found in the high RI group were stomach adenocarcinoma (STAD) (8.9% RAS mutated), bladder urothelial carcinoma (BLCA) (8.3%), head and neck squamous cell carcinoma (HNSC) (5.8%), cervical squamous cell carcinoma and endocervical adenocarcinoma (CESC) (5.2%), lung squamous cell carcinoma (LUSC) (3.5%) and oesophageal carcinoma (ESCA) (1.2%). RAS is not significantly mutated in these cohorts when compared to those with a lower RAS mutation ratio (Supplementary figure 5b). In order to explain the presence of these cancers in the highly RAS active group, we looked at the correlation between RAS pathway alteration status and mean RI across the cohorts. We defined RAS pathway alteration status as the number of patients with at least one alteration in a RAS pathway gene (as defined in TCGA driver pathway analysis^19^) leading to pathway activation. We identified a pan-cancer correlation (spearman coefficient 0.432, p-value 0.0135) (Figure 6c). Stomach (STAD), bladder (BLCA), head and neck squamous cell (HNSC), oesophageal (ESCA) and lung squamous cell (LUSC) cancers all had RAS pathway alteration rates above 50% and fell within the 99% confidence interval suggesting RAS pathway alterations other than RAS are driving activity in these cancers. The high RI values in endocervical (CESC) remain unexplained since the high frequency of *PIK3CA* mutation (28.5%) in this cancer is not significantly associated with RI (Supplementary figure 5c).

**Figure 6.**
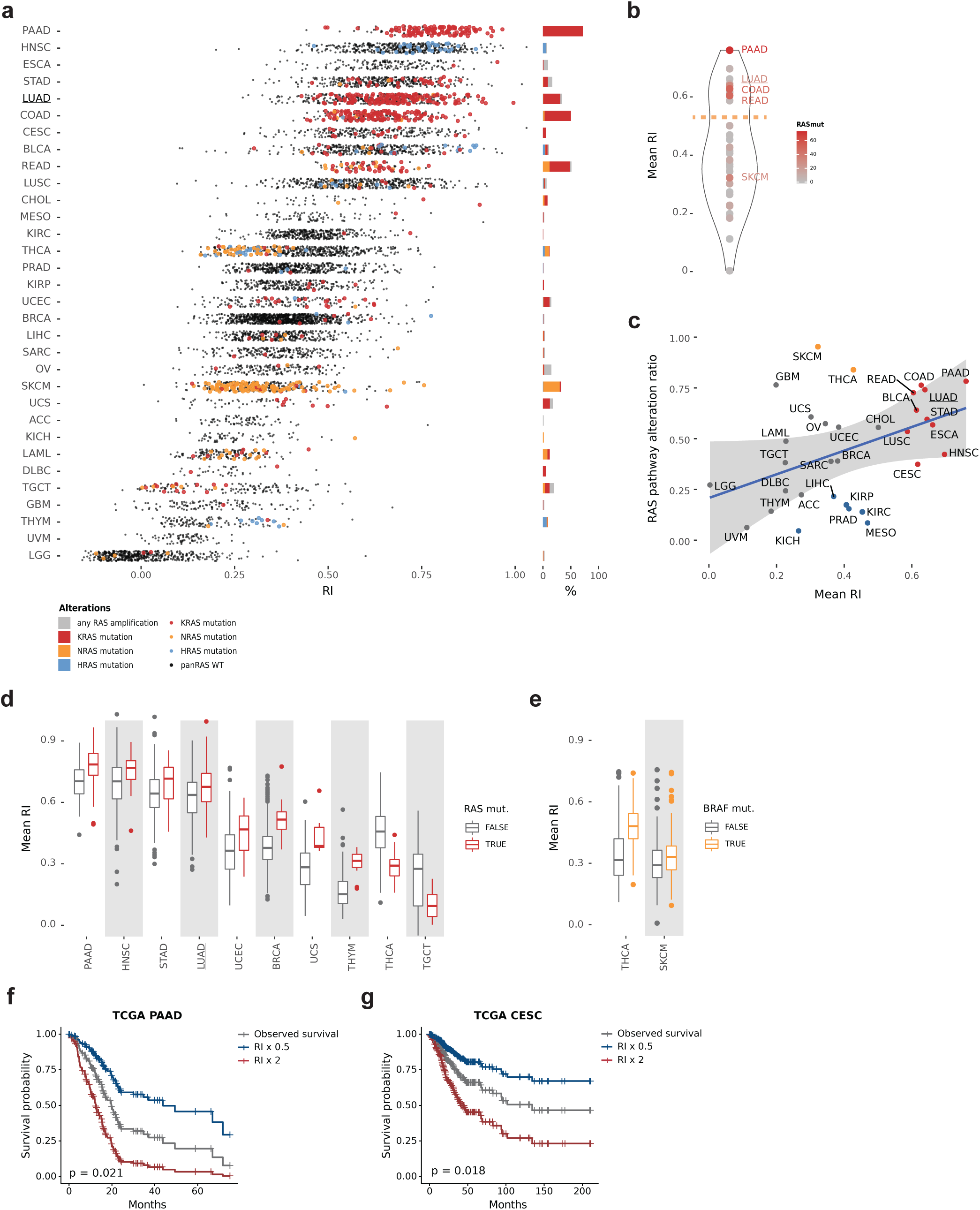
RAS Activity Pan-cancer. **(a)** RI plotted per-patient across the TCGA pan-cancer cohort. RNA-Seq gene counts were VST and z-score normalised per cohort. RAS mutants are highlighted (KRAS in red, HRAS in blue and NRAS in orange). Relative RAS isoform mutation frequencies per-cohort are shown in the barchart to the right using the same colours. Frequencies of patient with a RAS gene amplification but no mutation are shown in grey. **(b)** A violin plot showing the distribution of mean RI across each TCGA cohort. The dotted orange line indicates the distribution minima separating the two cancer populations. **(c)** The ratio of patients with one or more RAS pathway mutations plotted against the mean RI for each cohort. A linear regression fit is shown in blue with a 99% confidence interval shown by the grey ribbon (spearman coefficient 0.432, p-value 0.014). Highly RAS active tumours are shown in red, BRAF-driven tumours in orange and tumours with a RI value below the lower 99% CI are shown in blue. **(d)** Boxplots showing distributions of RI values for pan-RAS mutant and wild type patients. Significant cohorts are shown (linear model fit fdr < 0.05). **(e)** Boxplots showing distributions of RI values for THCA and SKCM split by BRAF mutation status (linear model fit fdr < 0.05). **(f)** Kaplan-Meier plot showing a Cox proportional-hazards regression fit of TCGA PAAD cohort survival data to corresponding RI values. The grey survival curve shows the observed data. The blue survival cure is the predicted survival if the RI values decreased 2-fold. The red curve are the predicted survival values if the RI were to increase 2-fold (coxph p-value 0.21). **(g)** Kaplan-Meier plot showing a Cox proportional-hazards regression fit of TCGA CESC cohort survival data to corresponding RI values. The grey survival curve shows the observed data. The blue survival cure is the predicted survival if the RI values decreased 2-fold. The red curve are the predicted survival values if the RI were to increase 2-fold (coxph p-value 0.018).

Skin cutaneous melanoma (SKCM) and thyroid carcinoma (THCA) mean RI were lower than predicted by their RAS pathway alteration ratio (Figure 4c, indicated in orange). Interestingly, *NRAS* is the main mutated isoform of RAS in these two cancers. However, RAS mutation does not correlate with RI in SKCM (Supplementary figure 5d) and shows a significant inverse correlation in THCA (Figure 6d) suggesting RAS is not the main driver of RAS activity in these cancers. Interestingly THCA and SKCM have the highest proportion of BRAF mutation (57% and 50%) compared with the next most common, colon (10%). We found BRAF mutation to be significantly associated with high RI values (Wilcox p-value THCA <2e-16, SKCM 2e-3) (Figure 6e) suggesting BRAF to be a key driver of oncogenic RAS activity in these two cancers. Given the lower than expected mean RI values it is possible that BRAF activation does not capture the full complexity of RAS pathway activation, possibly due to the use of multiple effector enzyme families by RAS proteins.

We measured a moderate RI in mesothelioma (MESO), kidney renal clear cell carcinoma (KIRC), prostate adenocarcinoma (PRAD), kidney renal papillary cell carcinoma (KIRP), kidney chromophobe (KICH) and liver hepatocellular carcinoma (LIHC) (Figure 6c, indicated in blue). All fall below the lower 99% CI indicating they have higher than expected mean RI values given their RAS pathway alteration ratio. The absence of RAS pathway alterations suggests that these cancers activate RAS via other mechanisms than the RAS pathway alterations considered here.

To further validate that RAS84 expression could predict high RAS activity in RAS mutants in individual cancers, we looked at pan-RAS mutation (*KRAS*, *NRAS*, *HRAS*) distributions across RI values per cohort (Supplementary Figure 5d). We identified a significant positive association between RAS mutations and RI in pancreatic (PAAD), LUAD, head and neck squamous (HNSC), thymoma (THYM), breast invasive carcinoma (BRCA), uterine carcinosarcoma (UCS), and uterine corpus endometrial carcinoma (UCEC) (Wilcoxon fdr<0.05) (Figure 6d, Supplementary table 9), reinforcing the idea that RAS84 can predict RAS activity within these cancers. For two cancers, thyroid (THCA) and testicular germ cell tumours (TGCT), we observed a negative association of RAS mutation with RI. As mentioned previously, THCA has high *BRAF* mutation levels with some *NRAS* but few *KRAS* mutations. TGCT is characterised by low levels of both *KRAS* (8%) and *NRAS* (3%) mutations. The higher RI activity in the RAS wild type group suggests that there may be a strong activator of RAS signalling in these samples, or a negative feedback loop in the mutants that is not apparent from the mutational data.

To determine if RAS84 had prognostic qualities in cancers other than LUAD, we ran Cox proportional-hazards regression analyses of overall survival against RI values on cohorts within our highly RAS active cancer group. We were able to identify a prognostic association with RAS84 expression for CESC and PAAD (p-value <0.05). Overall survival data with predicted outcome given a two-fold increase and decease in RI are shown in the survival plots (Figure 6f-g). This result shows RAS activity to be a possible prognostic indicator in other RAS-driven (pancreatic - PAAD) and RAS pathway active (cervical - CESC) cancers.

## Discussion

The importance of mutations in *RAS* oncogenes in tumourigenesis, cancer progression and resistance to treatment has been demonstrated in numerous model systems *in vitro* and *in vivo*. However, the mutational status of *KRAS* has neither prognostic nor predictive value in human lung cancer, limiting the possibilities to anticipate patient survival and to adapt treatments. From this, one might be tempted to conclude that lung adenocarcinomas lacking a *KRAS* mutation have acquired functionally similar patterns of signalling network activation to those with *KRAS* mutations. However, the development here of a transcriptional measure of oncogenic RAS activity has allowed us to distinguish significant differences in outcome and response to chemotherapy between lung adenocarcinoma patients. We also show that a high proportion of *KRAS* wild type tumours do exhibit the characteristics of RAS pathway activation. This analysis will facilitate the study of the effect of RAS on survival, cancer progression and resistance to treatment in patients and could ultimately inform clinical decisions.

Several groups previously developed “RAS-addiction” or “MEK-sensitivity” signatures using RNA interference against RAS, MEK inhibitor or farnesyltransferase inhibitor (when RAS-specific inhibitors were not available) to investigate resistance to RAS pathway targeted therapy^23, 41–43^. Others have also reported approaches to assess RAS activity in tumours from expression data. Bild and colleagues mapped several transformation signatures to expression data from lung tumours and identified a population of patients with deregulation in RAS, Src, β-catenin and Myc activity and poor survival^24^. Sweet-Cordero and colleagues assessed the enrichment of a *Kras*^V12^ tumour signature derived from a mouse cancer model in human tumours, observing an enrichment of the signature in human lung adenocarcinoma, but not specifically in KRAS mutants^44^. Nagy and colleagues reported a prognostic value in NSCLC using the mean expression of the top 5 deregulated genes in KRAS mutants versus non-mutants to segregate patients^45^. Way and colleagues used a machine learning approach and trained their classifier to detect *KRAS, HRAS* and *NRAS* mutations and copy number variation across cancers using the TCGA pan-cancer dataset ^46^. Of the five published RAS signatures we tested, just two predicted outcome in univariate survival analysis^24, 41^, but only RAS84 conserved its prognostic value when corrected for tumour stage in the same cohort. This finding suggests we were able to extract the RAS-target-genes which capture RAS activity and not only tumour aggressiveness.

We chose to derive our meta-signature in lung cancer because lung adenocarcinoma is known to be RAS-driven with a 30% *KRAS* mutation rate, but also has 26% patients that are RAS pathway wild type (no genetic alterations on any of the broader RAS pathway members), thus, presenting a potentially wide RAS-activity dynamic range. Our method offers an alternative approach to previously published methods in that it does not rely on the initial segregation of RAS mutant and wild type patients. We started from genes expressed in RAS active conditions, and we identified those that were good markers of KRAS mutants compared with non-activated RAS pathway tumours. This approach makes our method also sensitive to RAS active tumours driven by non-KRAS mutations. This nuance lacked in previously published attempts and could explain the greater performance of our approach over others to capture the aggressiveness induced by oncogenic RAS activity. Moreover, using cell line data in the initial derivation of the signature provided a pure tumour cohort, free from the complexities of the tumour micro-environment, which could have introduced noise in identifying the driver genes, thus demonstrated by the low expression of RAS84 in stromal and immune cells.

Our analysis shows that 84% of lung adenocarcinomas exhibit clear evidence of RAS pathway activation independent of *KRAS* mutation status. We show that RAS-pathway-mutation burden is associated with RAS84 activity in our pan-cancer analysis, demonstrating the influence of other RAS-pathway-member mutations. However, there are undoubtedly other indirect mechanisms driving RAS oncogenic activity in KRAS wt tumours, such as epigenetic regulation, inter-exonic variants, influence of the tumour microenvironment and growth factor expression, negative feedback loop regulation, metabolic regulation or others. Using a transcriptional approach presents the advantage of being agnostic to all upstream regulation and negates the complete characterisation of all drivers affecting RAS signalling. We identified four groups with different degree of RAS84 activity. The coincident mutations we observed in RAG-1 to -4 have previously been associated with specific phenotypes by Skoulidis *et al*. in a cohort of KRAS mutant lung cancer^30^. Our classification includes *KRAS* wt patients (representing 70% of all LUAD), thus broadening the clinical benefits of stratification to all patients.

Many studies over the years described the role of mutant *KRAS* in cancer progression, which might be expected to affect patient survival^5^. Oncogenic RAS promotes cell proliferation^47–52^, suppresses apoptosis^53^, shifts the metabolic program of cancer cells to sustain hyperproliferation^54, 55^, promotes angiogenesis^56^, increases inflammation^57, 58^ and remodels the extracellular matrix^59, 60^. Moreover, RAS promotes immune evasion by impairing antigen presentation^61^, recruiting immunosuppressive cells^62, 63^ and inducing immune checkpoint ligand expression^10^. Despite extensive literature describing how oncogenic RAS increases tumour aggressiveness, these findings do not reflect patients’ survival or response to treatment. An analysis of 227 patients with surgically resected NSCLC showed no association between RAS mutation and relapse^64^; a meta-analyse of 29 studies had shown no relation between RAS mutation and survival in lung cancer^65^. The discrepancy between laboratory experiments and observations made in the clinic could be explained by the fact that some *KRAS* wt tumours can still activate the RAS pathway due to events other than RAS oncogene mutations, such as *BRAF* or *EGFR* mutation. Multiple factors can affect RAS activity from one tumour to another in human cancer, unlike in the controlled isogenic systems typically used in laboratory studies where the only perturbation is the RAS mutation. Based on transcriptional activity, our approach is thus at least partially agnostic to a precise position in the signalling network of genomic alterations, explaining its superiority to predict outcome. Additionally, RAS84 expression appears broadly clonal, showing that RAS activity is generally an early driver event in lung adenocarcinoma. This observation is an essential consideration for developing clinical biomarkers when assessing tumour RAS activity from a single biopsy.

Adjuvant therapy is currently the first-line treatment for patients with early-stage lung cancers^66^. Although KRAS mutant promotes resistance to chemotherapy in isogenic experiments *in vitro* and *in vivo*^2–5^, it has no predictive value in patients with lung cancer^6, 7^. Using an independent cohort of lung adenocarcinoma (TEMPUS), we show that RAG-3 and -4 patients have a worse progression-free survival in response to first-line chemotherapy. This result is supported by the resistance to 23 chemotherapy drugs we observed in RAS-high lung cell lines *in vitro*. In both analyses, the classification based solely on the mutational status of KRAS did not reveal increased resistance to chemotherapy in KRAS-mutant cell lines or tumours. We also compared the response to drugs in cell lines mutated on any RAS pathway members versus all RAS pathway wt. Interestingly, this classification was not sufficient to predict resistance to chemotherapy, suggesting that other events –perhaps non-genetic– also affect the RAS oncogenic activity we capture in our approach. Surprisingly, RAG-2 showed the best response to chemotherapy. The absence of coincident mutations on tumour suppressor genes may explain the better response observed in this group than other RAGs.

We also evaluated the RAS activity signature across different cancer types. In a pan-cancer analysis across all cancer types, we demonstrated high signature expression in tumours known to be RAS-driven, and we showed RAS84 expression to be predictive of RAS pathway mutation burden across cancer types. We predicted nine cancer types to be highly RAS pathway active, almost all of which had a high representation of mutations in RAS pathway genes. Five cancer types (liver hepatocellular carcinoma, kidney renal clear cell and papillary carcinomas, prostate adenocarcinoma and mesothelioma) with low RAS pathway mutation burden classified as moderate RAS pathway activity, indicating that events other than genomic alterations activate RAS signalling in these cancers. In addition, the relationship between RAS84 expression and RAS pathway gene mutations is different in cancers with very high *BRAF* mutation levels, such as thyroid carcinoma and melanoma, which score as only moderate for RAS pathway activity. This suggests that the oncogenic RAS pathway’s output varies depending on the mutated gene, with the *BRAF* mutation not being equivalent to *KRAS* mutations. This might be expected from our knowledge of the bifurcating nature of the RAS pathway with multiple effector enzyme families directly targeted by RAS proteins. Our results show a great variability of RAS activity amplitude across cancers, highlighting the importance of assessing RAS activity per cancer cohort. Interestingly, the correlation between RAS84 expression and overall survival in pancreatic cancer, where 95% of tumours are mutated on *KRAS*, shows the direct link between RAS transcriptional activity and tumour aggressiveness.

Our finding that RAS84 correlates with resistance to chemotherapy needs to be validated in a larger cohort with more early-stage patients. Predicting response to chemotherapy in these patients is crucial to inform clinical decisions regarding the benefit of adjuvant therapies. It would be interesting to evaluate whether RAS84 predicts response to other treatments such as targeted therapy and immunotherapy. Defining the predictive ability of RAS84 for immunotherapy would be particularly relevant since we and others have shown that mutant KRAS protein can modulate the expression of immuno-suppressive proteins^9, 10^. Skoulidis and colleagues showed that KRAS and STK11/LKB1 co-occurring mutations are associated with poor response to PD-1 blockade in NSCLC patients^67^. In our classification, RAG-1 is enriched in STK11/LKB1 mutants, suggesting that this tumour group could be refractory to immune checkpoint blockade (ICB). In the same study, the authors showed that KRASm; TP53m tumours responded better to anti-PD-L1. RAG-3, and to a lesser extend RAG-4, are enriched in TP53 mutants, suggesting that these tumours could respond to ICB. About 20% of patients with NSCLC respond to ICB. The predictive value of RAS84 classification in response to ICB should be investigated in a following-up study.

The current large number of genes in RAS84 is a limitation to translating our classifier to a clinical assay. We kept the original 84 identified genes to capture the complexity in RAS-target expression adequately. However, it is conceivable to reduce the number of these genes by identifying key RAG-classification drivers and develop an assay that would be more suited to the clinic.

RAS84 captures RAS oncogenic activity in tumour samples better than the mutational status of *KRAS* when applied to cohorts of lung adenocarcinoma patients and other cancer types. We believe that the stratification of patients based on RAS84 expression will facilitate the study of the effect of RAS on survival, cancer progression and resistance to treatment in patients and could ultimately help clinical decision making.

## Methods

### Selection of the founder gene sets

We selected gene sets from several published data: the RAS addiction signature contained 380 genes upregulated in 5 KRAS-dependent cell lines (4 lung cell lines and 1 pancreatic cell line) compared with 5 KRAS-independent cell lines (4 lung cell lines and 1 pancreatic cell line)^23^; the KrasLA signature contained 89 upregulated genes in mouse lung tumours induced by the spontaneous recombination of KrasLA2 allele compared with normal lung and expressed in human lung tumours^44^; the HRAS transformation signature contained 245 genes correlating with the classification of HMEC samples into oncogene-activated/deregulated versus control^24^; the RAS pathway signature contained 105 genes previously curated from 3 studies including HRAS transformation, KrasLA and a signature of Salirasinib-treated human cancer cell lines^42, 43^; the MSigDB signature is the HALLMARK_KRAS_SIGNALING_UP meta-signature from MSigDB, which contained a list of 200 genes identified from overlaps between KRAS-related gene sets in other MSigDB collections^25^. We also generated a gene set from in-house data. The data was previously generated using the colon cancer cell line HCT116, which carries a KRASG13D mutation, and its isogenic cell lines Hke3 and Hkh2 where the *KRAS*^G13D^ allele was deleted by homologous recombination^68^. An Affymetrix analysis was performed on parental and recombined cell lines, and on sh-*KRAS* and sh-control in the HCT116 cell line. *KRAS*^G13D68^ was derived by selecting the genes upregulated (L2FC>1.5) when *KRAS*^G13D^ was expressed in all three experiments: HCT116 sh-*KRAS* versus control (6 days), Hke3 versus HCT116 and Hkh2 versus HCT116. We also identified a number of other oncogenic expression signatures to use as controls throughout the analysis (Supplementary table 10).

### CCLE

The CCLE microarray expression and mutation data were obtained from the CCLE legacy repository hosted at The Broad (https://data.broadinstitute.org/ccle_legacy_data). We selected lung-derived cell line data. We labelled the cell lines as either *KRAS* mutation positive, RAS pathway mutation positive or RAS pathway mutation negative. Cell lines were labelled as RAS pathway mutation negative if they had no mutation in a RAS pathway gene member defined in Sanchez-Vega and colleagues^19^. We removed cell lines with mutations in RAS pathway members other than *KRAS* from further analysis selecting 166 cell lines. We filtered the RAS signatures for genes relevant in the context of lung cancer. We modelled the log COV of expression against the mean log2 expression across all genes using loess regression. Signature genes with positive residuals with respect to the fit and a log2 mean expression value > 6 were selected.

We mapped each of the signatures to the cell line expression data and clustered the cell-lines using hierarchical clustering with a Ward.D2 agglomeration method. We split the resultant dendrograms into three clusters, labelling each high, low and unclassified for signature expression. The labels were assigned based on ranked mean cluster expression. To assess the performance of each of the signatures we calculated the significance of association of high and low clusters with *KRAS* mutation using a chisq test. To further refine each of the top 3 performing signatures, we ran a differential analysis between the high and low clusters using the Limma package from Bioconductor. We selected signature genes with an fdr < 0.05 as those driving the clustering. We merged the differential genes for each signature tested to form our meta-signature, RAS84. We identified RAS-high dependent transcriptional changes between RAS-high and RAS-low groups by limma (3.40.2) analysis on CCLE RMA normalised intensity estimates. Genes with a RAS group mean intensity estimate < 6 in both groups were removed from the analysis. Differential genes were selected by FDR < 0.05 and absolute LFC > 1. We ran the GO analysis using the clusterProfiler (3.12.0) package from Bioconductor testing all Biological Process terms from org.Hs.eg.db_(3.8.2) (FDR < 0.001)

### CCLE Drug Sensitivity Screen

We obtained drug sensitivity data from GDSC (IC50) (v1 367, v2 198 compounds)^69^ and CTRP (AUC) (v1 185, v2 481 compounds)^70^ for the CCLE cell lines. We clustered the VST normalised RAS84 CCLE RNA-Seq data into two clusters, RAS high and RAS low. We tested for significant differences in drug response values across the two RAS activity clusters by linear model correcting for any KRAS mutation status effect (< 0.05 fdr). We analysed each of the two release versions separately for each of the two data repositories. We identified enriched compound target pathways in the GDSC results by hypergeometric test using the TARGET_CATEGORY annotation provided (< 0.05 fdr). We also tested for oncogenic KRAS mutant dependent and oncogenic KRAS pathway dependent drug responses in the GDSC data. We used genotype data from the CCLE. We called the RAS pathway as mutated if any of the pathway genes contained a mutation^19^.

### Patient Data sets

#### TCGA Pancancer Data

All TCGA RNA-Seq gene-level read-counts were downloaded using the TCGAbiolinks (TCGAbiolinks_2.8.4) package from Bioconductor ^71^ (legacy=TRUE). Raw counts were VST normalized using the varianceStabilizingTransformation function within DESeq2 (DESeq2_1.20.0) from Bioconductor^72^. Normal samples were removed prior to analysis. To compare across the cancer cohorts, we z-score normalized samples and genes. Mutation data were obtained from Sanchez-Vega and colleagues^19^ and specific *KRAS* mutation data was downloaded from TCGAbiolinks and integrated with the expression data. Survival and proliferation data were obtained from Thorsson and colleagues^73^. RAS84 gene annotations were mapped to the RNA-Seq feature ids (Supplementary table 2).

#### Seoul lung adenocarcinoma cohort, GSE40419

RNA-Seq RPKM values for 87 adenocarcinoma patients were downloaded from GEO using the getGEO function from the GEOQuery Bioconductor package. RPKM values were log2 transformed prior to cluster analysis. Mutation data were obtained from Seo and colleagues^74^. Where multiple features existed per-gene the one with the maximum mean expression value across the cohort was selected.

#### Uppsala II RNA-Seq

RNA-Seq gene-level read-counts and clinical data were downloaded from the Gene Expression Omnibus (GEO GSE81089). The raw counts were VST normalised using the varianceStabilizingTransformation function within DESeq2 (DESeq2_1.20.0) from Bioconductor^72^. Ensembl gene annotations were obtained using the biomaRt package from Bioconductor. The 103 stage I, II, & III adenocarcinoma samples were selected prior to further analysis (column histology:ch1 == 2, stage.tnm.ch1 != 7). RAS84 gene annotations were mapped to the RNA-Seq feature ids (Supplement table 2). In the case of IER3, which maps to multiple features in this dataset, the feature with the largest mean VST value across all samples was selected (ENSG00000137331).

#### TRACERx

RNA-Seq gene-level read-counts were obtained for the TRACERx 100 patient cohort. The counts were VST normalized using the varianceStabilizingTransformation function within DESeq2 (DESeq2_1.20.0) from Bioconductor^72^. The 102 adenocarcinoma samples were selected (Histology == “Invasive adenocarcinoma”). The counts were further z-score normalised prior to SVM classification. RAS84 gene annotations were mapped to the RNA-Seq feature ids (Supplement table 2).

#### Lambrechts scRNA-Seq

Log2 CPM normalised scRNASeq data for B, T, fibroblasts, alveolar, EC and myeloid cells from five lung carcinomas were obtained from ArrayExpress (E-MTAB-6149).

#### TEMPUS CLINIC-GENOMICS

TEMPUS clinic-genomics is a retrospective lung cancer cohort Tempus clinic-genomic database containing 1,711 patients. Clinical data were extracted from the Tempus real-world oncology database of longitudinal structured and unstructured data from geographically diverse oncology practices, including integrated delivery networks, academic institutions, and community practices. All data were de-identified in accordance with the Health Insurance Portability and Accountability Act (HIPAA). The database extract was retrieved and de-identified in 2018 and contained cohorts with patients records spanning from 1990-2018. We identified 108 adenocarcinoma patients from the TEMPUS database with first line chemotherapy treatment and matched RNA-seq molecular data. We calculated a progression free survival interval from the associated patient clinical histories. We took the start time of treatment as time zero. We used a recorded recurrence event, a reported progressive disease outcome, a progression in reported tumour stage, death or the administration of an alternative therapy as an end-point to progression free interval. In the cases of repeated chemotherapy treatment, we took a gap between treatments of > 6 months as a PFS end point. In the absences of any endpoint events we censored on the last follow up time if no neoplasm was recorded or the last reported outcome if it was one of stable disease, progression free or partial response. We also integrated stage, the administration of radio therapy, age and sex data. We applied VST and z-score normalisation to the RNA-Seq gene level counts across all 633 adenocarcinoma patients in the cohort. We classified patients into RAG groups using our SVM classifier.

### RAG Classification

The RAS84 TCGA LUAD VST expression matrix was clustered using hierarchical clustering with a Euclidian distance measure and a ward.D2 agglomeration method (hclust function, R). We split the dendrogram into five clusters and labelled them RAG-0 to RAG-4 based on their mean RAS84 expression value across all samples, lowest to highest. We also calculated an RI value for each sample, defined as the mean VST value across the RAS84 genes. We repeated this analysis for each of the original founder signatures. We assessed the performance of each of the signatures by testing for statistically significant differences in the observed *KRAS* mutation frequencies across the five groups using a chi-squared test (chisq.test function, R). We tested all somatic variants reported in Sanchez-Vega and colleagues^19^ (N>10) for significant frequency differences across the RAS84 clusters using a chi-square test (fdr < 0.05). Mosaic plots were generated using the vcd package from R (vcd_1.4-4). Specific *KRAS* mutation genotypes were tested individually against a background of all remaining samples using a chi-squared test. The Seoul cohort (GSE40419) was clustered in the same way as the TCGA samples. KRAS, EGFR and TP53 mutations were tested for significant differences in observed frequencies across the five RAGs using a chi-squared test. To identify RAG driver genes, we identified genes with the largest deviation in expression from the mean across all samples. We first calculated RAG mean expression values per gene. We then scaled these values to the mean across all samples to calculate a RAG deviation value. Genes with an absolute deviation of > 1 were selected. These genes were clustered across RAG mean values by hierarchical clustering using Pearson’s correlation and ward.D2 agglomeration (cor and hclust function, R). We identified RAG dependent transcriptional changes by comparing RAGs 1-4 to RAG-0 correcting for tumour purity in the model since we were interested in tumour specific effects. Tumour purity CPE values were obtained from^75^. Differential genes were identified using DESeq2 (1.24.0) (fdr<0.05) and shrunken LFC values were generated using the lfcShrink function with type=“ashr”^76^. The genes were further filtered using the shrunken LFC values prior to GO analysis (absolute shrunkenLFC > 1). Go analysis was carried out using goseq (1.36.0) from Bioconductor, accounting for the length bias inherent in RNA-Seq results. Only terms associated with ‘Biological Process’ were considered and enriched p-values were corrected using Benjamini & Hochberg correction (FDR< 0.05).

### RPPA

We obtained level 4 normalised TCPA LUAD RPPA data from https://tcpaportal.org/tcpa/download.html. We identified 349 samples in common between the TCPA LUAD RPPA cohort and our classified TCGA LUAD cohort. We found 216 assayed proteins with values across all samples. We fitted a linear model across the RAGs against RAG-0 as the control for each protein assay. A Benjamini & Hochberg FDR multiple testing correction was applied across all tests (lm and p.adjust functions from R).

### Lung adenocarcinoma OS and PFS analysis

We fitted a univariate Cox proportional-hazard model to RAG labels and TCGA LUAD overall survival data to test RAG as a predictor for outcome. We also fitted a univariate Cox proportional-hazard regression model using RI as a continuous predictor of outcome. We ran OS univariate and multivariate Cox proportional-hazard analysis against RAG and RI in the Uppsala cohort. Overall survival time was calculated by subtracting the surgery date from the vital date (columns vital.date.ch1 - surgery.date.ch1). We compared a reduced coxph model including TNM stage, World Health Organization (WHO) performance status, smoking history, gender and age covariates (columns stage.tnm.ch1, ps.who.ch1, smoking.ch1, gender.ch1 and age.ch1) to a full model including either RAG labels or RI values, using LRT with AVOVA. We accounted for possible non-linear age effects by applying a restricted cubic spline to age using the rcs function from the rms R package (rcs(age, 3 )). We performed a PFS multivariate coxph analysis using the TEMPUS patients. We constructed a reduced model using stage, radiotherapy, gender and age covariates applying a restricted cubic spline as above. We compared this model to a full model including RAG labels using LRT with anova. In our pancancer analysis we fitted univariate Cox proportional-hazard regression models to RI and overall survival data from each TCGA cancer cohort from our high RAS activity cancer group. We identified cancers where RI was a significant predictor of outcome (p-value < 0.05). Kaplan-Meier curves were produced as above. These analyses were carried out using the coxph function from the survival R package (survival_3.1-11). Kaplan-Meier curves were produced using the ggsurvplot function from the survminer R package (survminer_0.4.4). In the case of the RI analysis the Kaplan-Meier curves were generated using predicted survival data from the coxph model given a 2-fold increase or decrease in RI.

### Pan Cancer RAS84 analysis

To compare RAS84 expression pan-cancer we z-score normalized the previously VST normalised RNA-Seq TCGA data (see section: TCGA Pancancer Data) and merged across the 32 cohorts. To assess RAS activity per sample we calculated an RI value for each sample, defined as the mean expression across the RAS84 genes. To identify high and low RAS active tumours we plotted the distributions of the mean RI values per cohort. From the observed bimodal distribution of cancer RI mean values, we calculated kernel density estimates (density function, R) and split the tumours at the minima between the two population maxima (mean RI value 0.53) (Supplementary figure 5a). We identified variants enriched in the high RAS activity tumours by hypergeometic test (fdr < 0.05). To obtain a RAS pathway mutation view we calculated a RAS pathway mutation burden percentage for each sample. We labelled samples as being RAS pathway mutated if they had a mutation in any gene defined in the RAS pathway by Sanchez-Vega et al. We tested for a significant correlation between RAS pathway mutation burden and mean RI using a Pearson’s correlation test.

To test for an association between RI values and RAS mutation status, per cancer, we ran Wilcoxon tests across the RAS mutated and non-mutated groups. We merged the oncogenic mutation calls for KRAS, HRAS and NRAS prior to testing. Cancers with a total RAS mutation count < 5 were excluded from the analysis. False discovery rates were calculated to account for multiple testing. We identified significant cancers by FDR < 0.05.

### SVM classifier

To facilitate the RAS classification of lung adenocarcinoma RNA-Seq samples we constructed an SVM classifier. We used the RAG labels derived from the LUAD TCGA cluster analysis as class labels and the TCGA LUAD RAS84 expression matrix as training data. The raw gene-level counts were first VST and z-score normalized. We constructed a Radial Sigma SVM using the caret R package^72^. The train function was used to optimize the classifier using a cv resampling strategy with 10 iterations. The classifier was validated against a pre-selected test subset of patients.

### TRACERx classification

We classified the samples using the z-score VST RAS84 expression matrix and our SVM classifier constructed from the TCGA analysis results and detailed in the classifier methods section above. We called the presence SNVs in each of the groups (PhyloCCF score > 0.05). We calculated a sample distance matrix from the VST expression matrix using Euclidian distance. We plotted the density of distance measures for intra- and inter-tumour distances. We determined the degree of enrichment of RAS84 genes with stable intra-tumour expression, but high inter-tumour variance was assessed relative to the distributions of all expressed genes, in-line with the methods presented in Biswas and colleagues^33^.

## Acknowledgements

We thank Philippe Juin for helpful discussion at the beginning of this project, Crick Bioinformatics and Biostatistics group for their support, the Oncogene Biology Laboratory and Miriam Molina for helpful discussion and critical reading of the manuscript.

## Funding

This work was supported by funding to JD from the Francis Crick Institute, which receives its core funding from Cancer Research UK (FC001070), the UK Medical Research Council (FC001070) and the Wellcome Trust (FC001070), from the European Research Council Advanced Grant RASImmune and from a Wellcome Trust Senior Investigator Award 103799/Z/14/Z. SCT was funded in part by a Marie Skłodowska-Curie Individual Fellowship from the European Union (MSCA-IF-2015-EF-ST 703228 - iGEMMdev).

## Author Contributions

PE, SCT, CS and JD designed the study, interpreted the results and wrote the manuscript. PE and SCT performed the computational analyses. DB, MM and DH provided datasets for analysis. GK carried out statistical analysis. All authors contributed to manuscript revision and review.

## Competing interests

JD has acted as a consultant for AstraZeneca, Bayer, Novartis, TheRas, Vividion, Jubilant. CS is a co-founder of Achilles Therapeutics, receives grant support from Pfizer, AstraZeneca, BMS, Roche–Ventana and Boehringer Ingelheim, has consulted for Pfizer, Novartis, GlaxoSmithKline, MSD, BMS, Celgene, AstraZeneca, Illumina, Genentech, Roche–Ventana, GRAIL, Medicxi and the Sarah Cannon Research Institute, is a shareholder of ApoGen Biotechnologies, Epic Bioscience and GRAIL. TC and KS are employees of AstraZeneca. None of the other authors of this manuscript have a financial interest related to this work.

## Data and materials availability

The TRACERx tumour region gene-level RNA sequencing count data of 84 genes used during this study are available through the Cancer Research UK & University College London Cancer Trials Centre for non-commercial research purposes, and access will be granted upon review of a project proposal that will be evaluated by a TRACERx data access committee and entering into an appropriate data access agreement subject to any applicable ethical approvals. All other data associated with this study are present in the paper or supplementary materials, or as cited.

## Notes

### Competing Interest Statement

JD has acted as a consultant for AstraZeneca, Bayer, Novartis, TheRas, Vividion, Jubilant. CS is a cofounder of Achilles Therapeutics, receives grant support from Pfizer, AstraZeneca, BMS, Roche Ventana and Boehringer Ingelheim, has consulted for Pfizer, Novartis, GlaxoSmithKline, MSD, BMS, Celgene, AstraZeneca, Illumina, Genentech, Roche Ventana, GRAIL, Medicxi and the Sarah Cannon Research Institute, is a shareholder of ApoGen Biotechnologies, Epic Bioscience and GRAIL. TC and KS are employees of AstraZeneca. None of the other authors of this manuscript have a financial interest related to this work. 

