## Supplemental Figures 1-5 for "Oncogenic RAS activity predicts response to chemotherapy and outcome in lung adenocarcinoma"

Supplementary figure 1

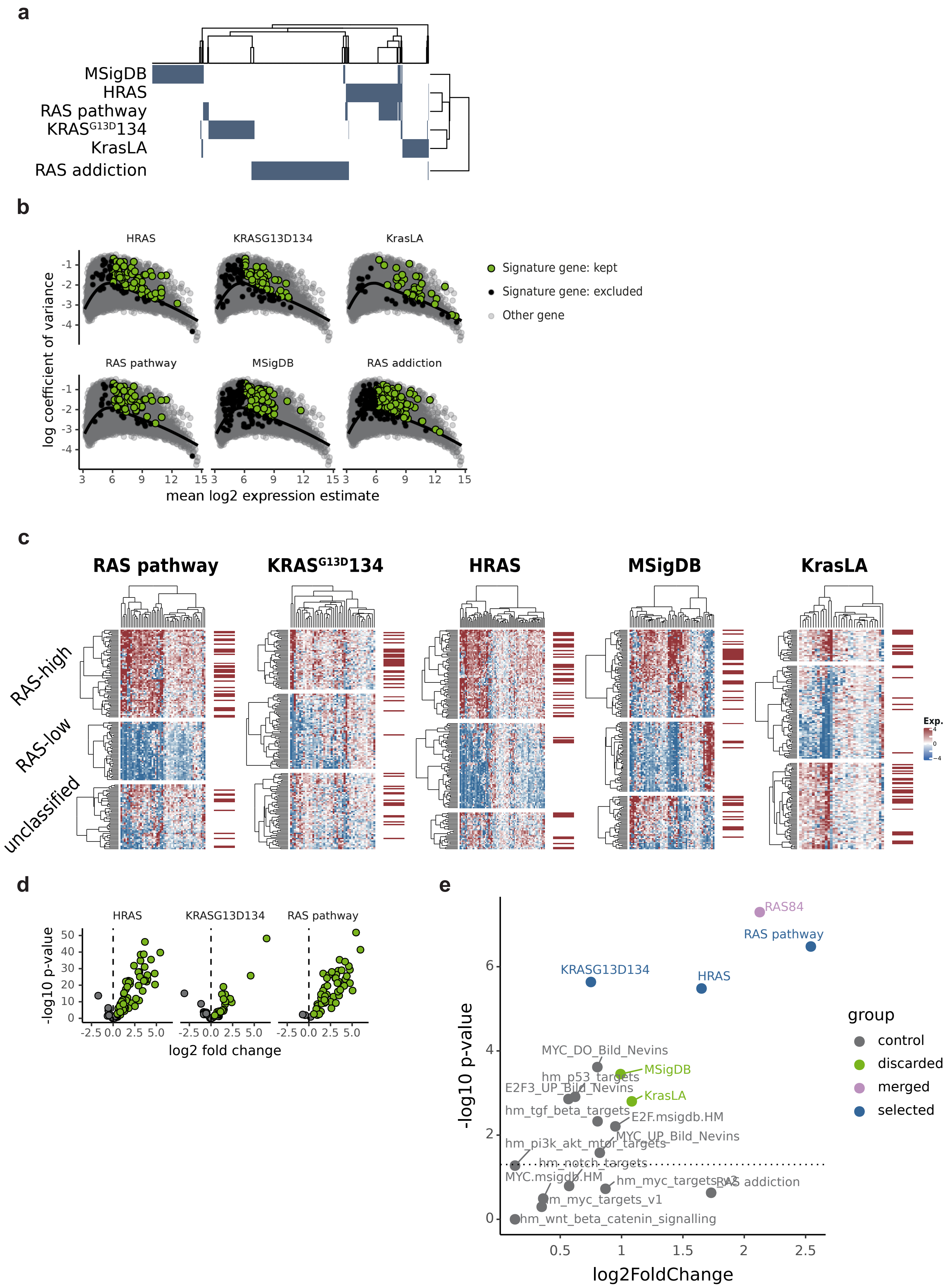

Supplementary figure 1

**Supplementary Figure 1. (a)** A heatmap showing the degree of overlap between the different RAS gene signatures. The genes are represented by the columns. Blue indicates the presence of a gene in a given signature. **(b)** Scatter plots highlighting RAS signature gene selection in the context of CCLE lung cell line expression data. Selected signature genes are highlighted in green, discarded in black and non-signature genes in grey. Log coefficient of variance was plotted against mean VST expression and a loess curve fitted to the data, shown by the black line. Signature genes with positive residuals with respect to the loess fit and a VST mean value > 6 were selected. **(c)** Heatmaps showing filtered RAS gene signatures (columns) across filtered (see methods) CCLE lung cell line expression data (rows). The expression of each gene across the cell lines is scaled to the median value. The cells and genes are clustered using hierarchical clustering with ward.D2 agglomeration. The cell lines are grouped by segmentation of the cluster dendrogram into three groups. The groups are labelled as RAS-high, RAS-low and unclassified based on the mean signature expression across the group. KRAS mutational status is indicated by the red bars to the right of the heatmaps. **(d)** Volcano plots showing the difference in expression between RAS-high and RAS-low groups for HRAS, KRASG13D134 and RAS pathway signatures. Genes with an fdr < 0.05 and a positive log2 fold change in the RAS-high group were selected, shown in green. **(e)** A volcano plot highlighting signature performance. The y-axis shows the significance of KRAS mutation segregation across RAS-high and RAS-low activity groups (-log10 p-value, chi-square). The x-axis shows the log2 fold change in mean expression between RAS-high and RAS-low activity groups. RAS84 meta signature is shown in purple, selected parent signatures in blue and discarded signatures are in green. A selection of RAS related controls gene signatures are shown in grey, highlighting the specificity of the selected RAS signatures to measure RAS activity.

### Supplementary figure 2

**a**

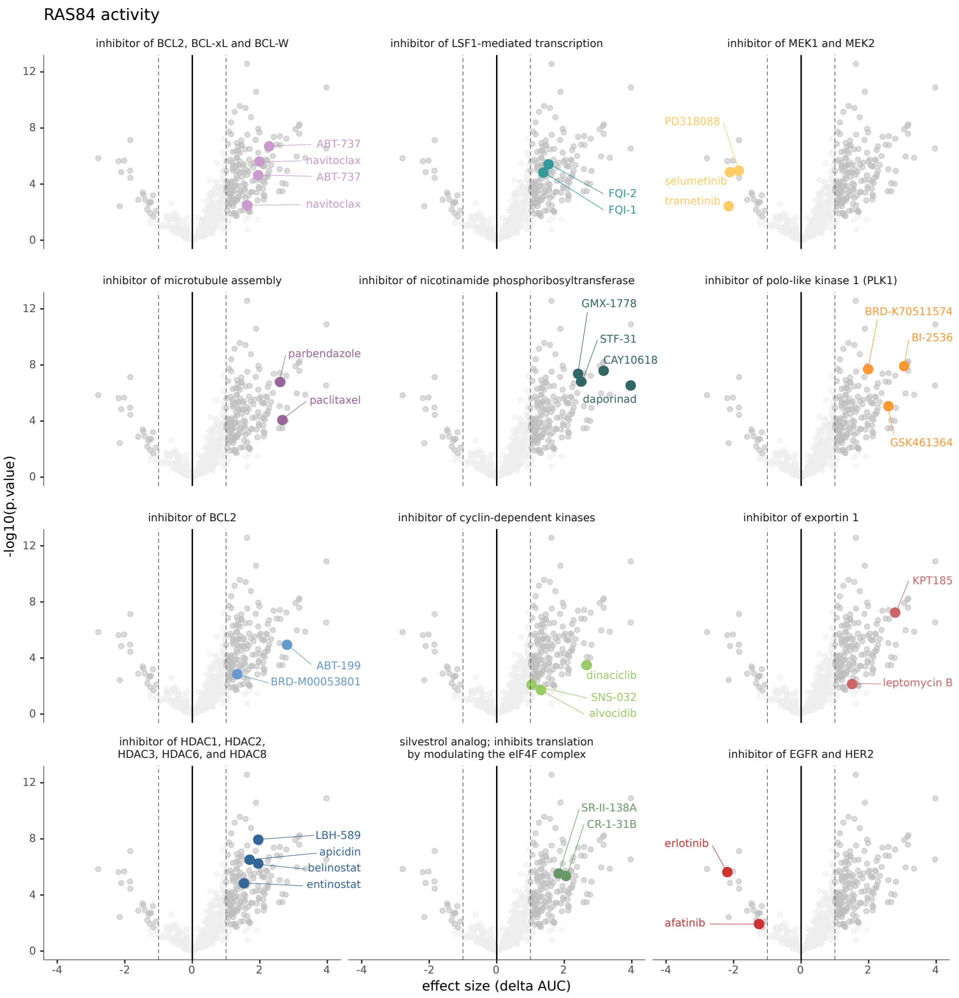


**Supplementary Figure 2.** **(a)** Volcano plots showing differences in IC50 values between RAS high and low CCLE cell lines. Drugs with enriched target annotations in the significant sensitive and resistant groups are highlighted. Drugs with an absolute log2 fold change > 1 and fdr < 0.05 are shown in dark grey. Results from both GDSC1 & 2 are shown.

Supplementary figure 3

**a**

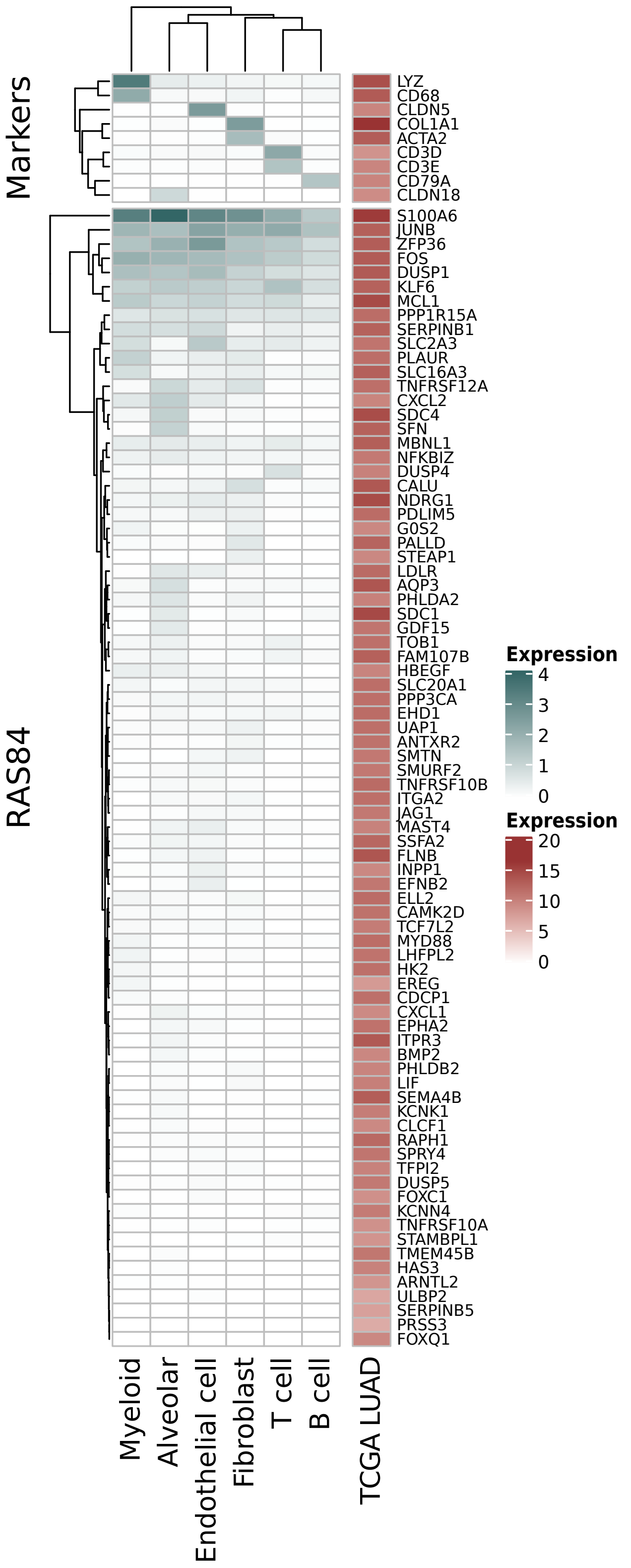

Supplementary figure 3

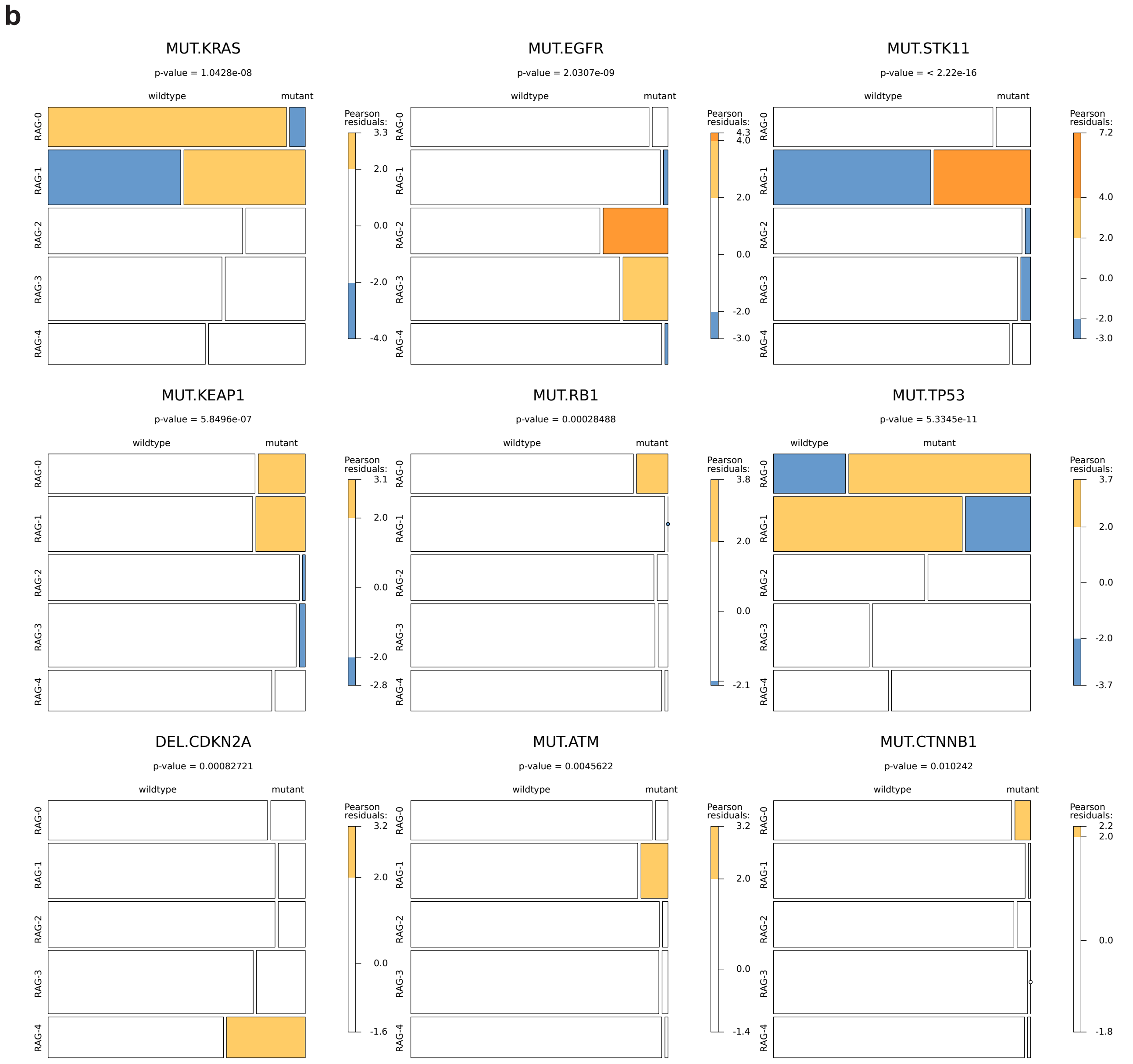


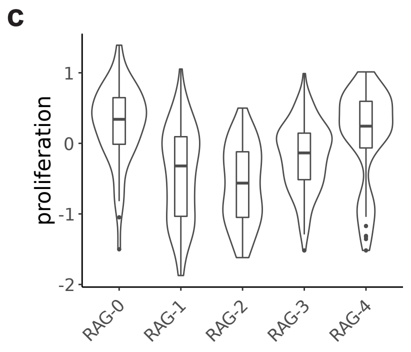


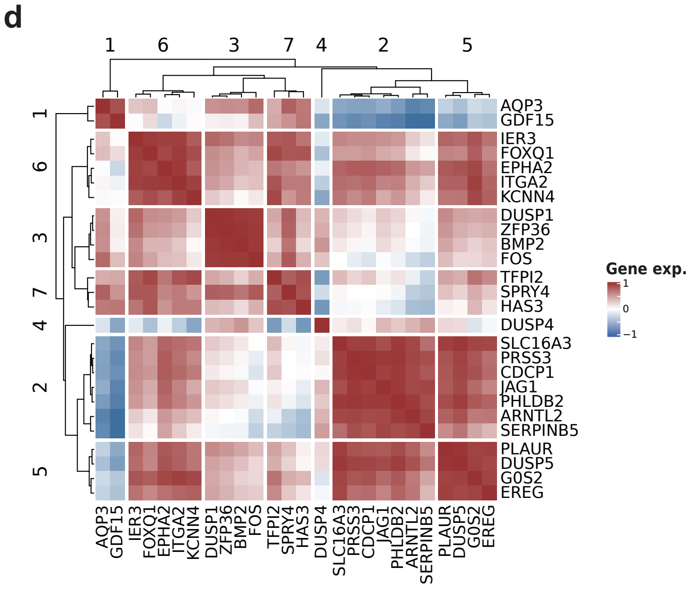

Supplementary figure 3

**Supplementary Figure 3. (a)** A heatmap showing minimal RAS84 expression detected in tumour infiltrating cells. The heatmap shows mean log2 CPM RAS84 expression across tumour infiltrating cell types from five NSCLC samples. The heatmap to the right shows mean VST normalised RAS84 expression across TCGA LUAD cohort. For reference a set of infiltrating cell expression markers are shown at the top. **(b)** Mosaic plots showing contingency table frequencies for given genetic variants across TCGA LUAD RAGs. The chisq test p-value is shown at the top of each plot. The plots were generated using the mosaic function from the vcd R package. The widths and heights of the rectangles represent the relative frequencies in each group. The colours are derived from the Pearson residuals and highlight over (orange) and under (blue) represented groups. **(c)** Distributions of TCGA LUAD sample proliferation scores per RAG. The proliferation score was taken from the TCGA pancancer data release. **(d)** Heatmap showing Pearson's correlation coefficients between RAS84 gene expression VST z-scores across the TCGA LUAD cohort. RAS84 genes with a deviation of > 1 log2 from the median VST value were selected. The genes were hierarchically clustered using ward. D2 agglomeration. The genes were grouped by segmentation of the dendrogram into seven correlated groups.

### Supplementary figure 4

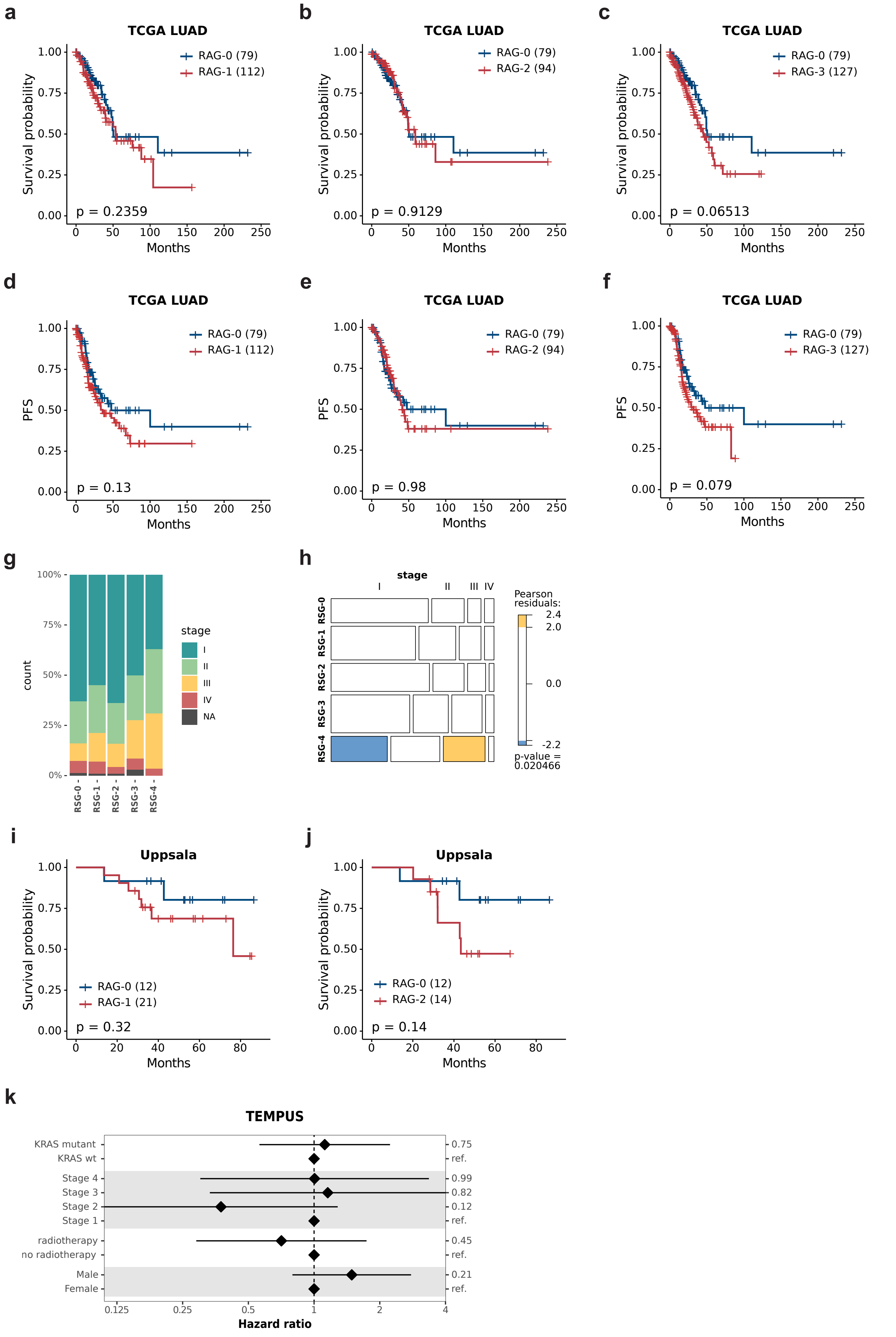

Supplementary figure 4

**Supplementary Figure 4. (a)** Kaplan-Meier plot showing overall survival data from the TCGA LUAD cohort for patients from RAG-1 and RAG-0 (coxph p-value 0.2359), the number of patients per group in indicated in brackets. **(b)** Kaplan-Meier plot showing overall survival data from the TCGA LUAD cohort for patients from RAG-2 and RAG-0 (coxph p-value 0.9129), the number of patients per group in indicated in brackets. **(c)** Kaplan-Meier plot showing overall survival data from the TCGA LUAD cohort for patients from RAG-3 and RAG-0 (coxph p-value 0.6513), the number of patients per group in indicated in brackets. **(d)** Kaplan-Meier plot showing progression-free data from the TCGA LUAD cohort for patients from RAG-1 and RAG-0 (coxph p-value 0.2359), the number of patients per group in indicated in brackets. **(e)** Kaplan-Meier plot showing progression-free data from the TCGA LUAD cohort for patients from RAG-2 and RAG-0 (coxph p-value 0.2359), the number of patients per group in indicated in brackets. **(f)** Kaplan-Meier plot showing progression-free data from the TCGA LUAD cohort for patients from RAG-3 and RAG-0 (coxph p-value 0.2359), the number of patients per group in indicated in brackets. **(g)** Stage percentages per RAG in TCGA LUAD. **(h)** Mosaic plot showing contingency table frequencies for TNM stage across TCGA LUAD RAGs with chisq test p-value. The widths and heights of the rectangles represent the relative frequencies in each group. The colours are derived from the Pearson residuals and highlight over (orange) and under (blue) represented groups. **(i)** Kaplan-Meier plot showing overall survival data from the Uppsala cohort for patients from RAG-1 and RAG-0 (multivariate coxph p-value 0.32), the number of patients per group in indicated in brackets. **(j)** Kaplan-Meier plot showing overall survival data from the Uppsala cohort for patients from RAG-2 and RAG-0 (multivariate coxph p-value 0.14), the number of patients per group in indicated in brackets. **(k)** Forest plot showing results from a multivariate Cox proportional-hazards analysis of PFS after chemotherapy in the TEMPUS lung adenocarcinoma cohort (n = 100 patients). KRAS mutants are compared to KRAS wt. Tumour stage, whether the patient received radiotherapy or not and sex were also tested (coxph p-value RAG-high p-value 0.0018). Hazard ratios and 5 and 95% confidence intervals are shown on a natural log scale.

Supplementary figure 5

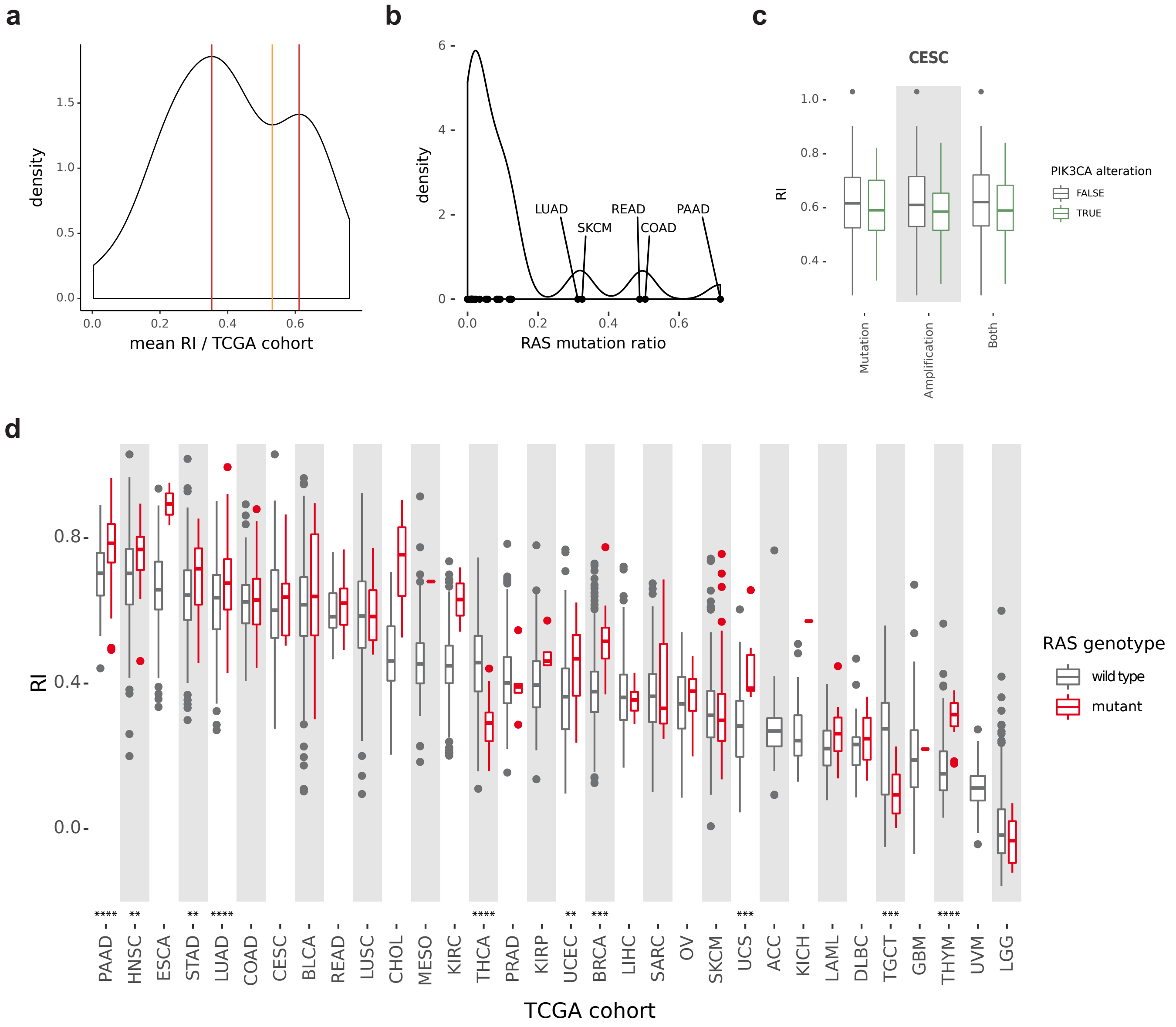


**Supplementary Figure 5. (a)** The distribution of mean RI values per cohort. The bimodal maxima are indicated by the red lines, the minima, segmenting the two populations of RAS activity cancers is indicated by the orange line. **(b)** The distribution of RAS (K, H and N) mutation frequencies in the different TCGA cohorts. **(c)** Boxplots showing RI distributions split by PIK3CA mutation, amplification and combined status for CESC. **(d)** Boxplots showing RI distributions per TCGA cohort split by RAS mutation status. Significant differences in RI means between the RAS mutant and wild type groups are indicated by the stars at the bottom of the plot (Wilcoxon, ****<0.0001, ***<0.001, **<0.01) (Supplementary table 6).
